# Secreted exosomes induce filopodia formation

**DOI:** 10.1101/2024.07.20.604139

**Authors:** Caitlin McAtee, Mikin Patel, Daisuke Hoshino, Bong Hwan Sung, Ariana von Lersner, Mingjian Shi, Nan Hyung Hong, Anna Young, Evan Krystofiak, Andries Zijlstra, Alissa M. Weaver

## Abstract

Filopodia are dynamic adhesive cytoskeletal structures that are critical for directional sensing, polarization, cell-cell adhesion, and migration of diverse cell types. Filopodia are also critical for neuronal synapse formation. While dynamic rearrangement of the actin cytoskeleton is known to be critical for filopodia biogenesis, little is known about the upstream extracellular signals. Here, we identify secreted exosomes as potent regulators of filopodia formation. Inhibition of exosome secretion inhibited the formation and stabilization of filopodia in both cancer cells and neurons and inhibited subsequent synapse formation by neurons. Rescue experiments with purified small and large extracellular vesicles (EVs) identified exosome-enriched small EVs (SEVs) as having potent filopodia-inducing activity. Proteomic analyses of cancer cell-derived SEVs identified the TGF-β family coreceptor endoglin as a key SEV-enriched cargo that regulates filopodia. Cancer cell endoglin levels also affected filopodia-dependent behaviors, including metastasis of cancer cells in chick embryos and 3D migration in collagen gels. As neurons do not express endoglin, we performed a second proteomics experiment to identify SEV cargoes regulated by endoglin that might promote filopodia in both cell types. We discovered a single SEV cargo that was altered in endoglin-KD cancer SEVs, the transmembrane protein Thrombospondin Type 1 Domain Containing 7A (THSD7A). We further found that both cancer cell and neuronal SEVs carry THSD7A and that add-back of purified THSD7A is sufficient to rescue filopodia defects of both endoglin-KD cancer cells and exosome-inhibited neurons. We also find that THSD7A induces filopodia formation through activation of the Rho GTPase, Cdc42. These findings suggest a new model for filopodia formation, triggered by exosomes carrying THSD7A.

## Introduction

Dynamic rearrangement of the actin cytoskeleton is critical for cell shape changes, elaboration of specialized cell structures, and cell movement. Thin actin-rich structures called filopodia are used for directional sensing and making initial contacts between neighboring cells ^1–3^. In migrating cells, filopodia protrude from the cell’s leading edge to promote cell polarization and facilitate adhesion ^4, 5^. Filopodia are also important for establishing cell contacts ^6–9^. In neurons, filopodia initiate synapse formation by contacting the axon of neighboring neurons and then develop into specialized postsynaptic structures called dendritic spines that are important for learning and memory ^10–13^. Filopodia have also been shown to contribute to invasive cancer cell behaviors, including metastasis ^4, 5, 14–21^.

Several different mechanisms have been described to promote filopodia formation, including direct nucleation of unbranched actin filaments by formins and other nucleators and reorganization of branched cortical actin filaments followed by actin polymerization ^1, 3, 22^. Additional actin regulatory proteins promote uncapping of the barbed ends, elongation and bundling of actin filaments. These actin regulatory proteins are controlled upstream by phosphatidylinositols, especially PI(4,5)P2, PI(3,4,5)P3, and PI(3,4)P2, and Rho GTPases, particularly Cdc42 ^1, 23^. Myosin-X may further promote filopodia maintenance by transporting and/or organizing cytoskeletal and adhesion molecules at filopodia tips ^6, 24–27^.

While the intracellular cytoskeletal machinery that drives filopodia formation has been heavily studied, there is little known about the extracellular cues that activate that machinery. Bone morphogenic proteins (BMPs) have been shown to regulate filopodia in endothelial cells, myoblasts, and neurons, often through activation of the Rho GTPase Cdc42 ^28–30^. VEGFA has also been shown to promote endothelial sprouting and filopodia formation ^31^. In neurons, a variety of extracellular molecules have been shown to regulate filopodia formation and dynamics, including brain-derived neurotrophic factor ^32^, Slit ^33^, Netrin-1 ^34^, and nerve growth factor ^35^. Filopodia formation has also been shown to be influenced in tumor cells by TGF-β signaling-induced alterations in gene expression, specifically upregulation of fascin protein ^20^. While these studies have identified some extracellular regulators of filopodia formation, the molecular mechanisms are poorly understood and some are indirect, via inducing gene expression changes in cytoskeletal components.

Exosomes are small extracellular vesicles (EVs) that are formed within multivesicular late endosomes (MVEs) and released into the extracellular space by fusion of MVEs with the plasma membrane. Exosomes carry bioactive cargoes, including proteins, lipids, and nucleic acids and promote autocrine and paracrine cell communication across a variety of systems ^36–38^. Notably, exosomes have been shown to promote polarization and motility of multiple cell types, including cancer cells, immune cells, and single-celled amoebae ^39–44^. Exosomes also play a key role in metastasis by seeding metastatic niches ^45–47^.

In previous studies, we found that exosome secretion is critical for the formation of two motility-related actin cytoskeletal structures: invadopodia and nascent cell-matrix adhesions ^42,48^. In our live imaging studies, we also frequently observed filopodia formation in close proximity to sites of exosome secretion and adhesion formation ^42^. Here, we directly investigate the role of exosomes in filopodia formation and stability. In both cancer cells and primary rat neurons, we find that genetic inhibition of exosome biogenesis or secretion inhibits filopodia formation and stability. In neurons, the reduction of filopodia was accompanied by a decrease in dendritic spines and synapses, which are structures that develop from filopodia. Filopodial defects of exosome-inhibited cells were fully rescued by the add-back of purified small EVs (SEVs, containing exosomes) but not larger ectosome-type EVs (LEVs). Proteomic comparison of SEVs and LEVs purified from melanoma cells identified the TGF-β co-receptor endoglin as a cargo that is unique to the SEV fraction. Endoglin was found to promote filopodia formation by cancer cells, as well as the filopodia-dependent behaviors of cancer cell metastasis and 3D motility. A quantitative proteomic comparison of exosomes purified from control and endoglin-KD melanoma cells revealed that the filopodia regulatory transmembrane molecule thrombospondin type 1 domain containing 7A (THSD7A) is reduced in endoglin-KD exosomes. Further investigation revealed that THSD7A is present on both cancer cell and neuronal exosomes and that recombinant purified THSD7A is sufficient to rescue filopodia defects resulting from exosome inhibition in both cell types. THSD7A-induced filopodia formation was diminished in the presence of a chemical inhibitor of the small GTPase Cdc42. Altogether, we find that exosomes drive filopodia formation in multiple cell types by carrying the filopodia regulator THSD7A.

Authors’ Note: Throughout this paper we use both the term “exosomes” and “small extracellular vesicles (SEVs)”. “Exosomes” is used when we have perturbed the endocytic pathway to specifically inhibit exosome biogenesis within MVE or secretion (via affecting docking of MVE with the plasma membrane) and can, therefore, attribute the biogenesis of those EVs to the endocytic pathway. “SEVs” is used for purified EV preparations where we have isolated a heterogeneous population of EVs and cannot define their origin.

## Results

### Exosome markers localize to filopodia

To visualize exosome association with cytoskeletal structures, we stained for CD63 in HT1080 fibrosarcoma cells, which form numerous filopodia. The cells were fixed and co-stained with phalloidin-Alexa Fluor 488 to visualize filamentous actin (F-actin) in filopodia. Confocal images of the cells show localization of CD63 puncta at the tips of filopodia (Fig. 1A, arrowheads). To observe the dynamic relationship between exosome secretion and filopodia formation, we utilized a live cell reporter that allows us to dynamically visualize the fusion of MVE with the plasma membrane, pHluorin-M153R-CD63-mScarlet ^49^. pHluorin is a pH-sensitive form of GFP that is non-fluorescent at acidic pH inside late endosomes and fluoresces at neutral pH ^50^, such as occurs upon endosome-plasma membrane fusion, whereas mScarlet is a pH-insensitive red fluorescent protein ^51^ and can report on MVE movements within the cell. Using this probe stably expressed in HT1080 fibrosarcoma cells, we observed localization of MVEs at (red signal) and fusion with (yellow puncta) the plasma membrane immediately before or coincident with filopodia initiation (Fig. 1B, Supplemental Movie 1). Quantitative analysis of the timing of yellow fluorescence appearance at the plasma membrane revealed that exosome secretion occurred with a median time of 20 seconds before filopodia formation (Fig. 1C,D). We also examined the localization of MVE markers to filopodia in primary cortical neurons by transiently transfecting neurons with the MVE docking factor GFP-Rab27b ^52^ along with mCherry as a cytoplasmic “filler” to visualize cellular neuronal structures. Notably, GFP-Rab27b localized to both the base and tips of neuronal filopodia (Fig. 1E,F), with more localized to the base than to the tips (63% vs. 37%, Fig. 1F). Together, both the cancer cell and neuron data suggest an association of MVEs and exosomes with filopodia.

**Figure 1.**
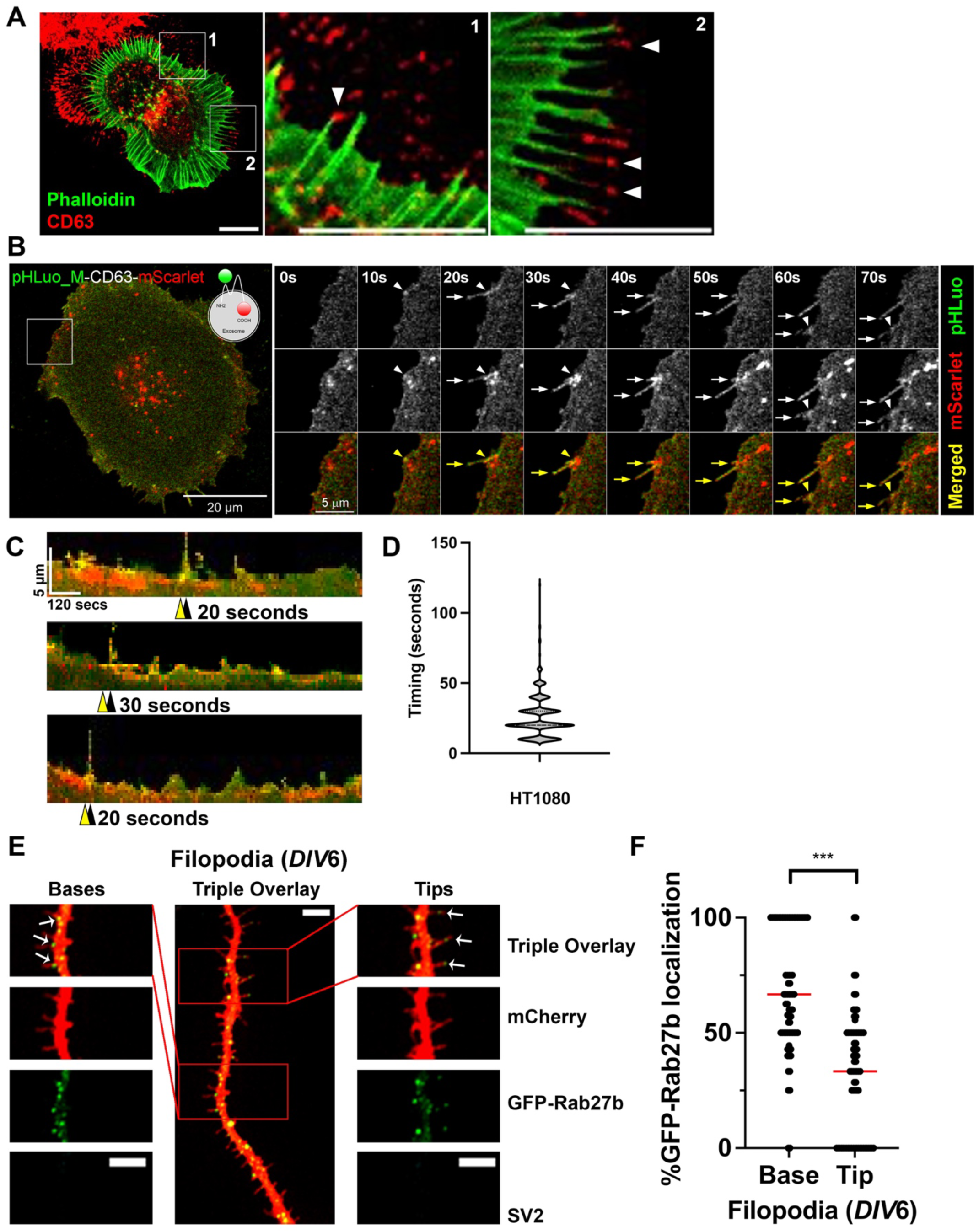
Exosome markers localize to the base and tips of filopodia in cancer cells and cortical neurons. A. Representative confocal image of HT1080 cells stained with phalloidin-Alexa fluor 488 and CD63 shown in red. The red channel has been edited using brightness and contrast tools for ease of visibility. Note localization of the exosome marker CD63 in extracellular deposits and at or near the tips of filopodia (arrowheads). Representative of 20 images. Scale bar is 10 μm in each panel. B. Time series of pHluorin-M153R-CD63-mScarlet movie in HT1080 cells. Yellow arrowheads indicate fusion sites and yellow arrows indicate filopodia. Note a filopodium forming shortly after MVE fusion. C. Representative kymographs showing MVE docking (red), fusion (yellow), and filopodia formation in HT1080 cells. Yellow arrowheads denote MVE fusion events and black arrowheads denote the formation of a filopodium. Each pixel is 10 seconds x 0.2857 μm. D. Quantification of the time elapsed between MVE fusion and filopodia formation. n=420 kymographs from 46 cells from 3 independent experiments. E. Primary cortical neurons were co-transfected with GFP-Rab27b (green) and mCherry as a filler to visualize filopodia (red) on *DIV*5 and fixed for imaging on *DIV*6. SV2 negative staining (no signal) identifies these structures as filopodia instead of dendritic spines. Arrows in merged images indicate localization of GFP-Rab27b to tips and bases of filopodia. Scale bars = 5 µm. F. Percent GFP-Rab27b localization to tips and bases of filopodia in 70 individual cortical neurons from three independent experiments. Red line indicates the median. Error bars, SEM. ns, not significant; * p<0.05; ** p<0.01; *** p<0.001

### Exosomes promote filopodia formation and stability in cancer cells

To directly test whether exosomes affect filopodia formation and/or stability, shRNA targeting Rab27a or Hrs was expressed in B16F1 melanoma cells (Fig. S1A,B), a frequently used cell line for studying filopodia, and shRNA targeting Rab27a was expressed in HT1080 cells (Fig. S2A). Rab27a is a key factor controlling MVE docking at the plasma membrane, which allows intraluminal vesicles to be secreted as exosomes ^52^. The ESCRT-0 protein Hrs plays an important role in the biogenesis of intraluminal vesicles within MVEs ^36, 53^. The exosome-enriched small EV (SEV) population was isolated by cushion density gradient ultracentrifugation ^54^, after isolation of large EVs by differential centrifugation from conditioned media (see methods). Consistent with current standards in the field ^55–57^, the EVs carried typical EV markers and were of the expected size and morphology by nanoparticle tracking analysis (NTA) and negative stain transmission electron microscopy (Fig. S1C-E and Fig. S2B,C).

Quantitation by NTA of EVs isolated from conditioned media confirmed that B16F1 Rab27a- and Hrs-knockdown (KD) cells secrete fewer SEVs compared to control cells (Fig. 2A). Consistent with a selective effect on MVE docking and biogenesis, these constructs had no effect on secretion of LEVs, which should contain primarily shed ectosomes (Fig. 2A). Phalloidin staining of actin filaments in Rab27a- and Hrs-KD B16F1 cells revealed reduced numbers of filopodia compared with control B16F1 cells (Fig. 2B,C). Similarly, inhibition of exosome secretion by Rab27a KD in HT1080 fibrosarcoma cells also led to a decrease in filopodia numbers (Fig. S2D-F). Consistent with a specific role for exosomes in controlling filopodia dynamics, the addition of SEVs purified from control B16F1 cells to Hrs-KD B16F1 cells rescued the defect in filopodia numbers (Fig. 2D,E). By contrast, there was no effect of LEVs on filopodia numbers. The addition of SEVs to B16F1 shScr control cells also increased filopodia number in a dose-dependent manner, while LEVs had no effect (Fig. 2F).

**Figure 2.**
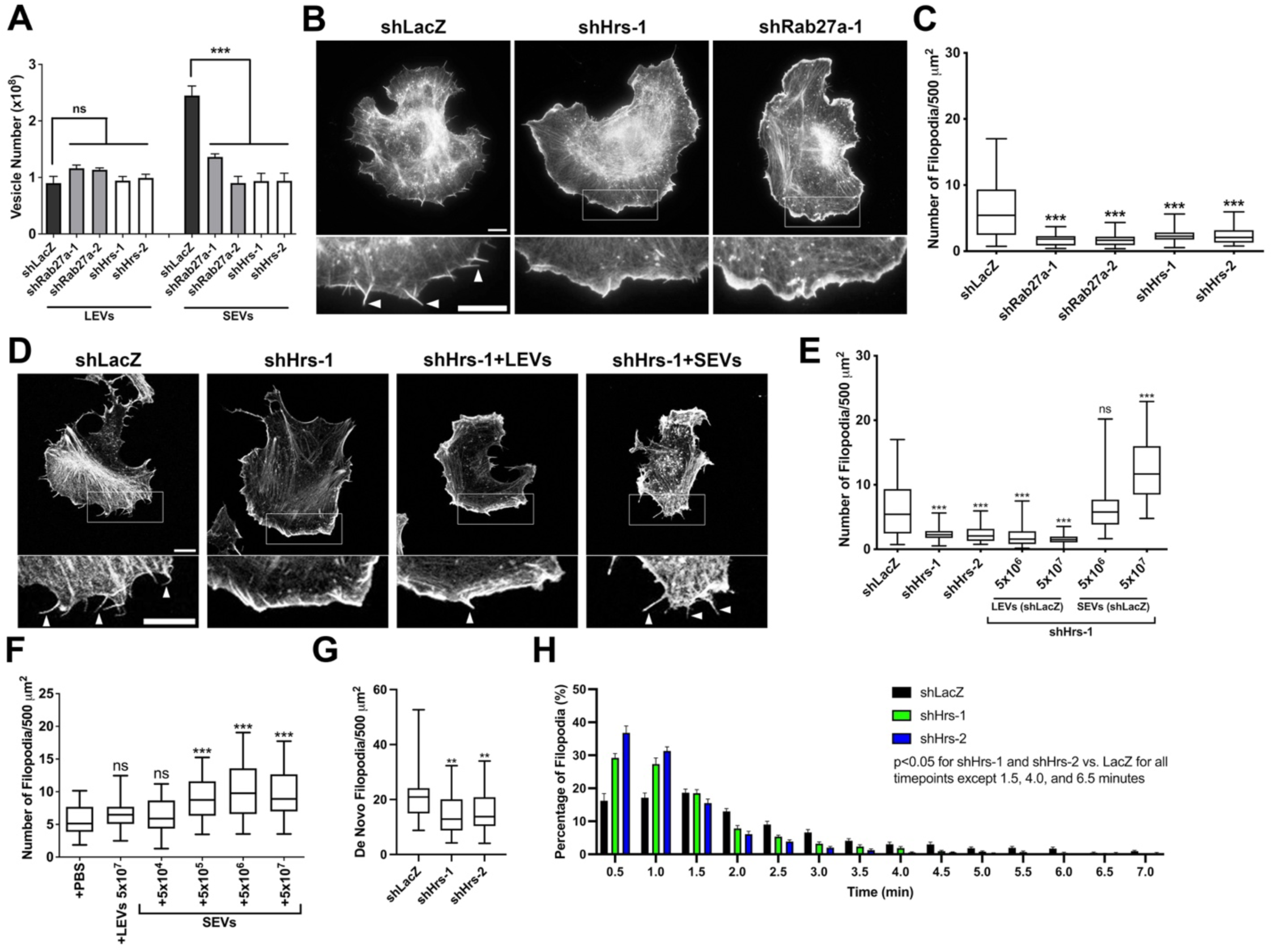
Exosomes promote filopodia formation and stability in cancer cells. A. EVs secreted from equal numbers of control (shLacZ), Rab27a-KD, and Hrs-KD B16F1 cells over 48 h were quantified using NanoSight particle tracking analysis (N=3). B. Representative images of control (shLacZ), Hrs-KD, and Rab27a-KD B16F1 cells stained with rhodamine-phalloidin. Arrowheads show examples of filopodia. Images have been edited with brightness and contrast for ease of visibility. C. Quantification of filopodia from images as in B (N=3, ≥ 27 total cells per condition). Filopodia number per 500 μm^2^ cell area. D. Representative images of filopodia in B16F1 control (shLacZ) and exosome-depleted (shHrs) cells treated for 18 h with LEVs or SEVs isolated from control cells. Arrowheads show examples of filopodia. Images have been edited with brightness and contrast for ease of visibility. E. Quantification of filopodia from images as in D (N=3, ≥ 20 total cells per condition). F. Filopodia number in B16F1 shScr cells treated with indicated numbers of purified LEVs or SEVs, for 18 hours (N=3, ≥25 total cells per condition). G. Control (shLacZ) and exosome-depleted (shHrs) B16F1 cells were transiently transfected with tdTomato-F-tractin to visualize filopodia formation. Live images were taken every 30 seconds for 15 minutes and newly formed filopodia were counted at each time point. Only filopodia that form and fully retract during the duration of the video were quantified. (N=3, ≥20 total cells per type) H. Lifetime of newly formed filopodia from (G). Lifetime is defined as the time from first formation of the filopodia to full retraction. Bars represent mean and error bars are SEM. (N=3, ≥20 total cells per type) Scale bars in wide field and zoom insets = 10 μm. Error bars, SEM. ns, not significant; * p<0.05; ** p<0.01; *** p<0.001.

To determine whether exosome secretion affects filopodia formation and/or stability, live imaging of B16F1 cells expressing the actin marker tdTomato-F-tractin ^58^ was performed. Analysis of the number of new filopodia formed over time, as well as the lifetime of those filopodia, revealed that Hrs-KD reduces both the formation (Fig. 2G) and stability (Fig. 2H) of filopodia (see also Supplemental Movie 2).

We quantified filopodia in tumor cells as “filopodia per cell area” to match with corresponding primary neuron data (Fig 3), in which we had to plot filopodia per 100 μm length to account for the fact that the entire neuron body was too large to fit in a field of view. To address concerns about this graphing choice, we have included the data presented in Figure 2 plotted as filopodia per cell (Fig. S2G-J). The data are highly similar, with the one exception being the de novo filopodia in shHrs-2 cells trended the same but was not statistically significant (Fig. S2J). It is possible that this data set is no longer significantly different than the control cells because the cell area of the shHrs-2 cells is slightly higher than that of control cells, although not significantly different.

**Figure 3.**
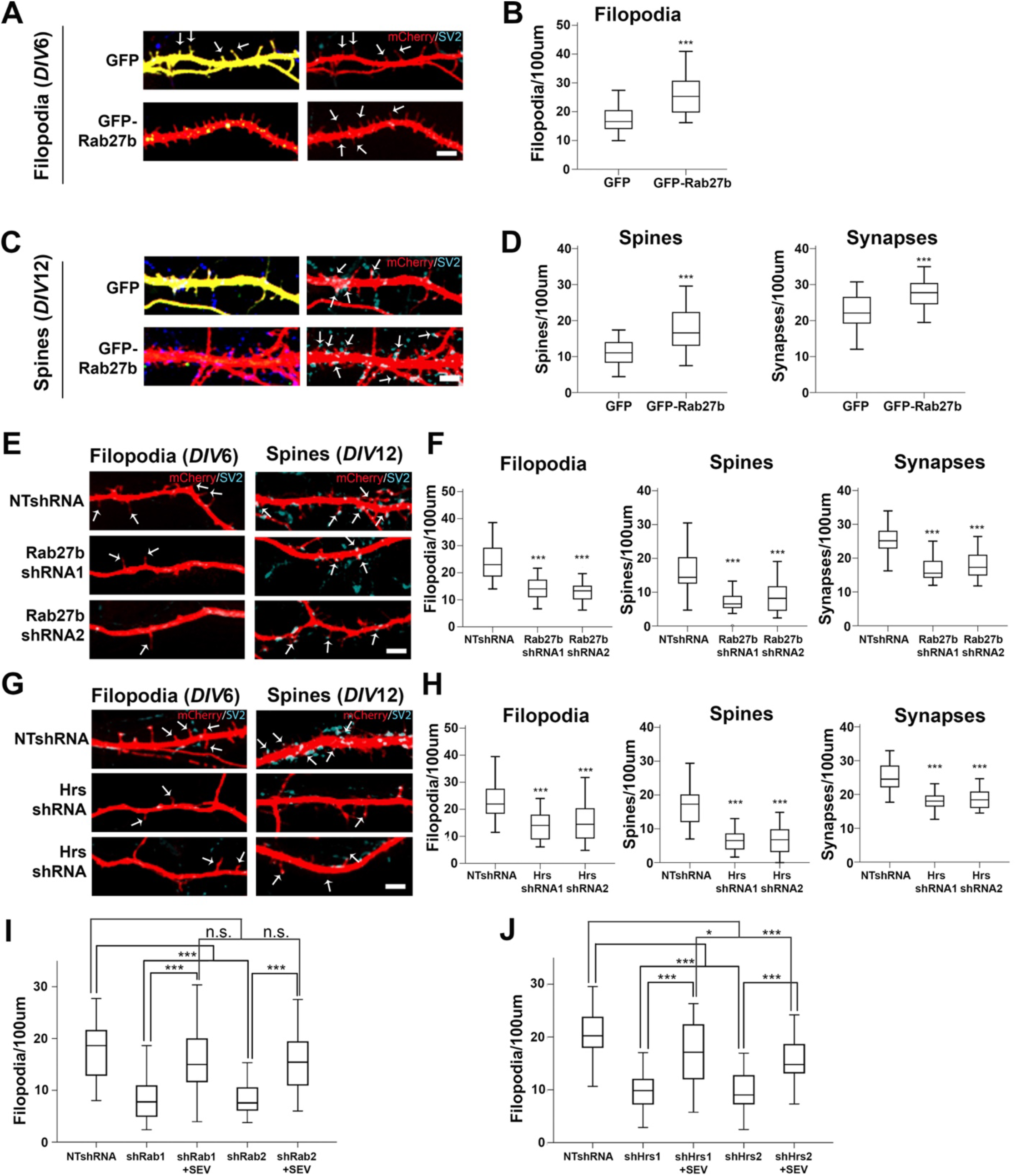
Exosomes promote filopodia, spine, and synapse formation in cortical neurons. Primary cortical neurons were co-transfected with plasmids for expression or inhibition of exosome regulatory molecules and mCherry (red) as a cytoplasmic filler to image neuronal protrusions, then fixed and immunostained with SV2 (pseudo-colored in cyan) to visualize synapses. Filopodia were identified as thin SV2-negative, mCherry-positive protrusions. Spines were identified as dendritic protrusions that co-localize with SV2. Synapses were identified as SV2-positive puncta present on both dendritic protrusions and dendritic shafts. A, C. Representative images of primary rat cortical neurons co-transfected at DIV6 when filopodia typically form (A) or DIV12 when synapses typically form (C) with GFP or GFP-Rab27b (green, left images) and mCherry (red) and co-stained with SV2 (blue, right images). B, D. Quantification of filopodia (DIV6), spine and synapse density (DIV12) from images as in A and C. E-G. Images and analysis from neurons transfected with control shRNA (NTshRNA) or shRNAs against Rab27b (E) or Hrs (G) and immunostained for SV2. Quantification of filopodia (DIV6), spine and synapse density (DIV12) for Rab27b-KD (F) or Hrs-KD (H) neurons. I, J. Rescue experiments. Control and KD neurons (as indicated) expressing shRNAs and mCherry were treated with purified SEVs on *DIV*5 for 24 hrs at a dose of 200 EVs per neuron, then fixed and stained for SV2 at *DIV*6. Filopodia, spine and synapse density were quantified from more than 30 primary or secondary dendritic shafts from three independent experiments for each condition. Scale bars = 5 μm. Error bars, SEM. *p<0.05, **p<0.01, ***p<0.001.

### Exosomes promote filopodia and synapse formation in neurons

In neurons, filopodia formation is critical for the subsequent development of synapses as filopodia mature into postsynaptic specializations called dendritic spines. In primary rat neuron cultures, numerous filopodia are present on dendritic shafts at day *in vitro* (*DIV)*6 while dendritic spines are predominant around *DIV*12. To test whether exosomes control filopodia in neurons, we altered expression of exosome regulators in primary rat hippocampal and cortical neurons. Both Rab27a and Rab27b regulate exosome secretion, by respectively promoting MVE docking with the PM and regulating the anterograde movement of MVEs to the PM ^52^. However, in some cell types, one form of Rab27 is expressed more highly than the other, suggesting some possible redundant functions ^59^. Since Rab27b is the predominant form of Rab27 protein present in the brain ^60, 61^ we overexpressed Rab27b in primary neurons. Cortical neurons overexpressing GFP-Rab27b exhibited a significant increase in the number of filopodia examined at *DIV*6, quantitated as thin protrusions that were negative for the synapse marker SV2 (Fig. 3A,B). This increase in filopodia density translated into an increased number of SV2-positive dendritic spines and synapses at DIV12 (Fig. 3C,D). A similar increase in filopodia, spine, and synapse density occurred in primary hippocampal neurons upon GFP-Rab27b expression (Fig. S3A,B).

To inhibit exosome secretion in primary cortical and hippocampal neurons, we knocked down Hrs or Rab27b by transient transfection of two different shRNAs along with mCerulean at least 48 hours prior to immunostaining. For both genes, shRNA-transfected neurons (recognized by mCerulean co-transfection) exhibited 40-60% reduction in protein expression analyzed by fluorescence intensity, compared to cells transfected with nontargeting shRNA (NTshRNA, Fig. S4A-D). Analysis of immunostained cells revealed that loss of either Rab27b or Hrs in KD neurons led to a significant reduction in filopodia, spines, and synapse density compared to NTshRNA controls (Fig. 3E-H, S3C-F). To further confirm that the effect of gene KD on filopodia density was due to loss of exosome release, we performed rescue experiments by treating Rab27b- or Hrs-KD primary cortical neurons for 24 hrs with SEVs isolated by differential centrifugation from *DIV*9 primary cortical neurons (Fig. S4E-G). LEVs were not tested due to the low recovery of LEVs from neuronal conditioned media. The dose of 200 SEVs per neuron was chosen based on the estimated SEV secretion rate from primary cortical neurons over the 24 h time period of the assay. In the Rab27b-KD condition, SEV treatment fully rescued the filopodia number defects of untreated KD controls (Figs. S4H and 3I). For Hrs-KD condition, there was partial rescue in filopodia density upon SEV treatment (Figs. S4I and 3J). These data indicate that - similar to cancer cells - endogenous exosomes secreted by neurons promote filopodia formation.

### Endoglin is an SEV cargo that promotes filopodia formation in cancer cells

EVs carry multiple bioactive protein cargoes. Because only SEVs but not LEVs were able to rescue or induce filopodia numbers in B16F1 melanoma cells (Fig. 2), we identified unique SEV cargoes by running SEV and LEV lysates on an SDS-PAGE gel, staining it with colloidal Coomassie, extracting 4 bands that were enriched in the SEV lysates (Fig. 4A, arrowheads) and performing proteomic analysis of trypsin digests of the extracted protein bands. Our approach was validated by the presence of typical exosomal/SEV proteins CD63, LAMP1, and the ESCRT-1 protein Multivesicular Body Subunit 12B (MVB12B) in one of the unique bands analyzed from the SEV lane (Fig. 4A, Supplemental Table 1). In SEV lane bands 1 and 2, we identified the TGF-β co-receptor endoglin (Fig. 4A). A full list of proteins identified in the SEV-enriched bands is shown in Supplemental Table 1. Western blot analysis confirmed that endoglin is enriched in SEVs compared to LEVs (Fig. 4B).

**Figure 4.**
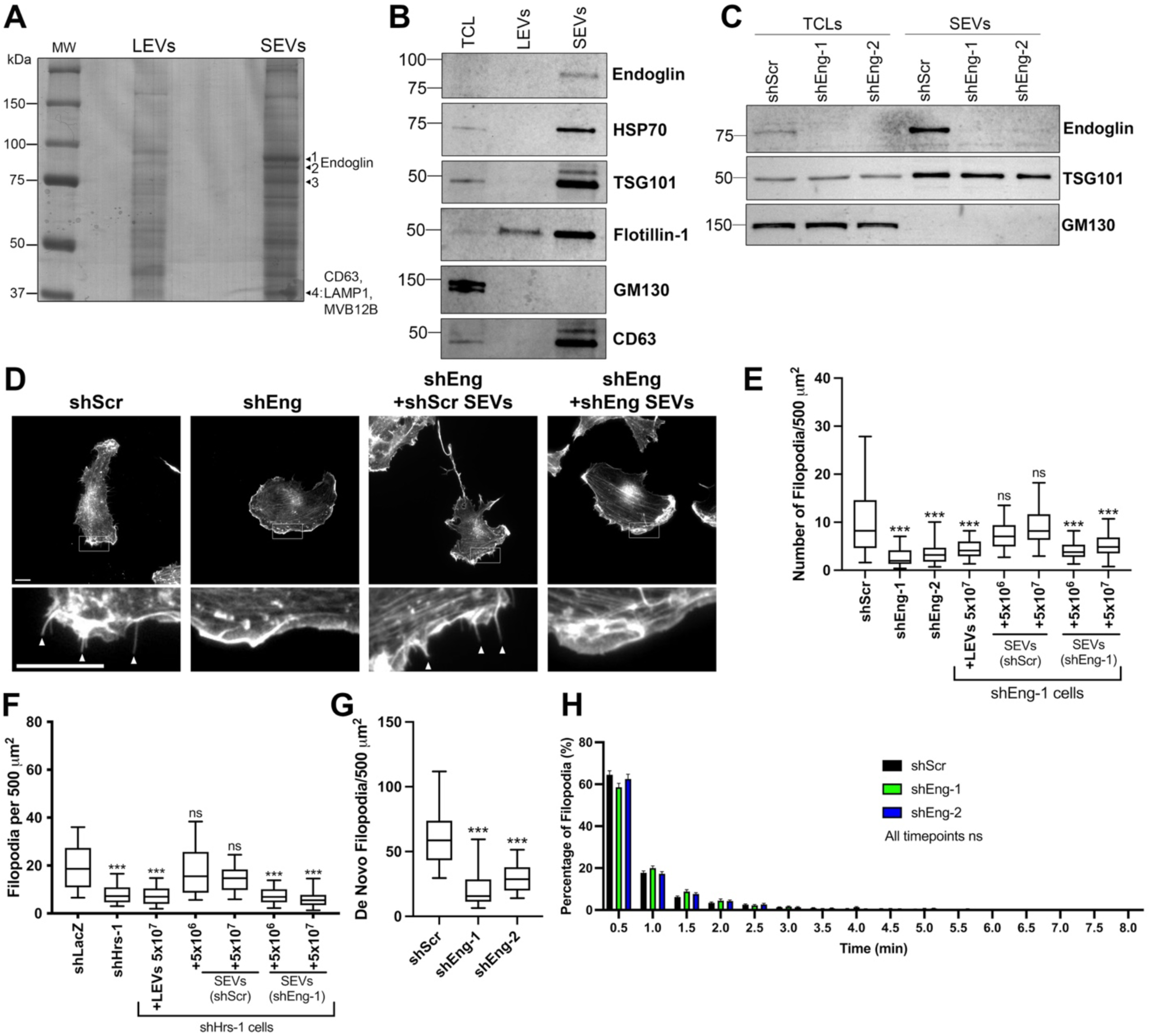
Endoglin is an SEV-enriched cargo that promotes filopodia formation. A. Purified LEVs and SEVs were run on a colloidal blue-stained gel. Four arrows denote SEV bands that were cut and submitted for proteomics, along with notable proteins identified (see Supplemental Table 1 for the full proteomics results). B. B16F1 total cell lysate (TCL), LEVs, and density gradient purified SEVs were run on an SDS-PAGE gel and probed by Western blot for endoglin, and EV positive (HSP70, TSG101, flotillin-1, and CD63) and negative (GM130) markers. C. Total cell lysate (TCL) and small EVs (SEVs) from endoglin-KD (shEng) and control (shScr) B16F1 cells were run on an SDS-PAGE gel and probed by Western blot for endoglin, EV marker TSG101, and EV negative marker GM130. D. Representative images from control (shScr) or endoglin-KD (shEng) B16F1 cell lines incubated for 18 h with no EVs (left panels), or with SEVs purified from control (+shScr SEVs) or shEng cell lines (+shEng SEVs) (right panels). Arrowheads indicate example filopodia. Scale bar = 10 μm. E. Quantification of filopodia in control (shScr) and endoglin knockdown (shEng) cells treated with the indicated number of LEVs or SEVs for 18 hours (N=3, at least 20 cells per condition per repetition). F. Filopodia number in B16F1 control (shLacZ) or exosome depleted (shHrs) cells treated with indicated numbers of LEVs, control (shScr) SEVs, or endoglin-KD (shEng1) SEVs for 18 hours. N=3, ≥ 20 cells per condition per rep. Representative images for this experiment are shown in figure S5E. G and H. B16F1 cells were transfected with tdTomato-F-Tractin and imaged live every 30 seconds for 15 minutes. Only filopodia that form and fully retract during the duration of each video were quantified. G. De novo filopodia formation. H. Filopodia lifetime, defined as the time from initial filopodia formation to full retraction. Bars represent mean and error bars are SEM. (N=3, β25 total cells per type) ns, not significant; * p<0.05; ** p<0.01; *** p<0.001.

Endoglin is a TGF-β co-receptor that regulates multiple processes relevant to motility and filopodia, including cellular signaling, adhesion, and cytoskeletal regulation ^62–65^. To test whether endoglin regulates filopodia formation, we stably expressed endoglin-targeting shRNAs in B16F1 melanoma cells and confirmed that endoglin levels are reduced in both SEVs and cells (Fig. 4C). Compared with control cells, stable endoglin-KD cells exhibit a reduction in filopodia number (Fig. 4D,E) but no significant effect on SEV release from cells (Fig. S5A,B). Likewise, transient KD of endoglin in B16F1 cells also leads to a significant reduction in filopodia numbers (Fig. S5C,D). Whereas the addition of LEVs to stable endoglin-KD cells has little effect on filopodia numbers, treatment with control SEVs fully rescues the filopodia number defects of endoglin-KD cells (Fig. 4D,E). Consistent with endoglin being a key SEV cargo that controls filopodia, add-back of endoglin-KD SEVs does not rescue the filopodia defects of endoglin-KD cells (Fig. 4D,E). Furthermore, control - but not endoglin-KD - SEVs also rescue filopodia defects of exosome-inhibited B16F1 Hrs-KD cells (Fig. 4F, Fig. S5E). To determine whether endoglin selectively controls filopodia formation or stability, we performed live imaging of endoglin-KD B16F1 cells. We found a specific defect of endoglin-KD cells in de novo filopodia formation (Fig. 4G) with no change in filopodia lifetime (Fig. 4H, see also Supplemental Movie 3). These data suggest that endoglin specifically affects filopodia formation whereas additional exosome cargoes may affect filopodia lifetime, since Hrs-KD cells had a defect in both filopodia formation and lifetime (Fig. 2G,H). Finally, we assessed the role of endoglin in filopodia formation in HT1080 fibrosarcoma cells. Similar to B16F1 cells, KD of endoglin in HT1080 cells reduced filopodia numbers (Fig. S6A-E). These data suggest that endoglin and/or endoglin-regulated cargoes carried by exosomes have important functions in controlling filopodia dynamics.

### Endoglin promotes cancer cell metastasis and cell motility

We previously observed that exosome secretion is critical for metastasis and directional migration of cancer cells in chick embryos ^42, 44^. Since filopodia have been implicated in both of these behaviors ^3, 4^, we tested the role of endoglin in promoting cancer metastasis in this model. The chick embryo chorioallantoic membrane (CAM) model has been established as an effective way to study experimental metastasis by intravenous injection of tumor cells ^66^. To seed tumor cells at metastatic sites, we injected fluorescently labeled control and endoglin KD HT1080 cells intravascularly. After 4 days, we harvested and imaged the chick CAM to visualize metastatic cells and colonies (Fig. 5A). Many cancer cells and colonies were present in the CAM from shScr control cells. By contrast, CAMs from chick embryos injected with endoglin-KD HT1080 cells had reduced numbers of individual metastatic cancer cells and colonies (Fig. 5B-D).

**Figure 5.**
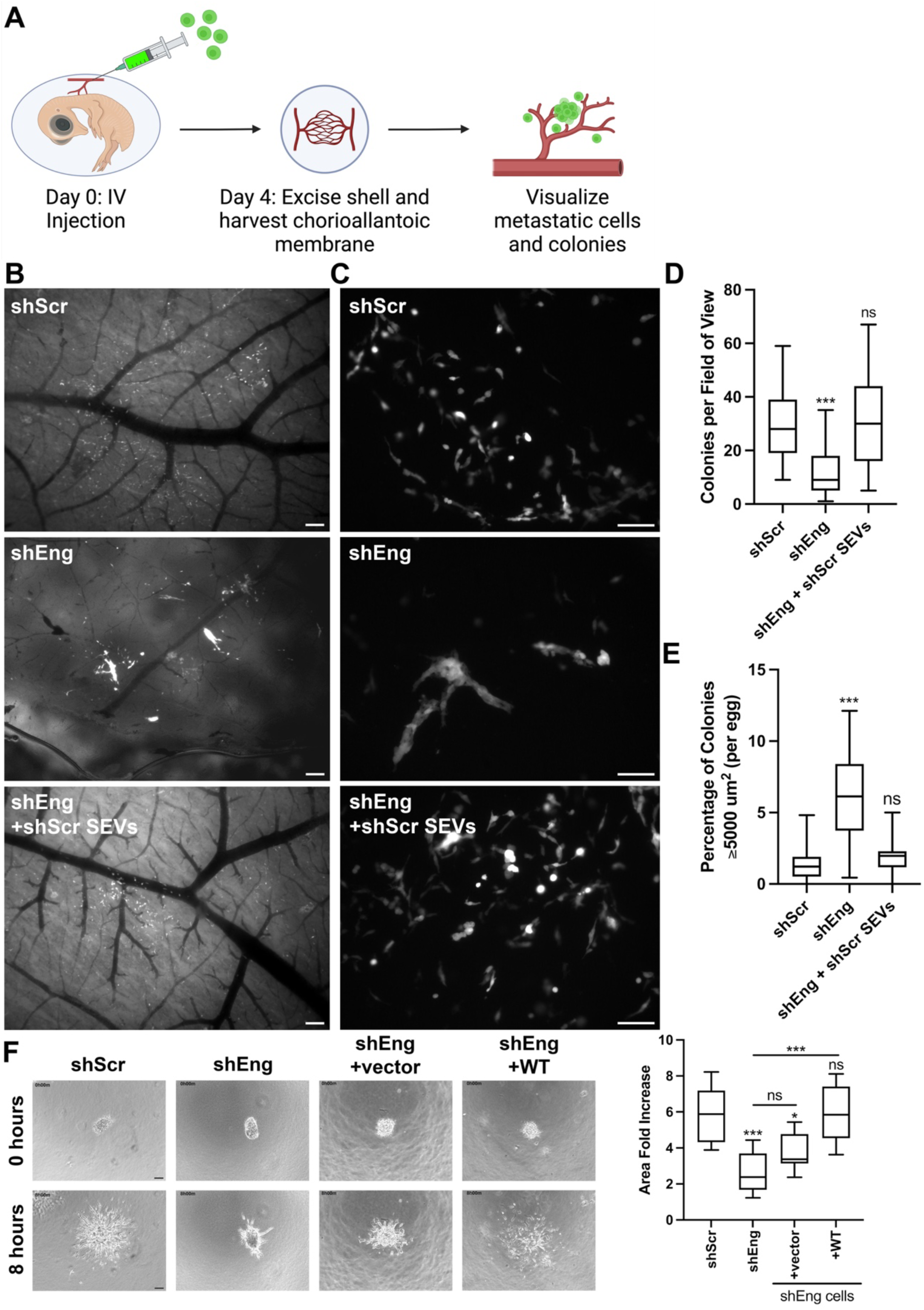
Exosomal endoglin controls motility and metastatic colony formation. A. Cartoon diagram of metastatic colony assay in avian embryos. On day 0, fluorescent HT1080 cells were injected (100,000 cells per egg) into the vein of the chicken embryo. On day 4, the egg was opened, the embryo was sacrificed, and a circular tool was used to punch holes through the shell. The chorioallantoic membrane (CAM) was peeled away from the shell, placed on a glass slide with a coverslip, and immediately imaged. The cartoon was created using BioRender. B. Representative low power wide field images of colony formation in the CAM. Scale bar = 200 μm. C. Representative high power wide field images of colony formation in the CAM. Scale bar = 100 μm. D and E. Quantification of CAM colony number (D) and size (E) from high power images as in C. 4-7 eggs were harvested per condition. D. Colony number is graphed per field of view using 25-30 fields of view per egg. E. Quantification of the percent of large (≥ 5000 μm^2^) colonies formed by control and shEng HT1080 cells. F. 3D invasion in collagen. HT1080 cell spheroids were seeded in collagen gels and imaged for 8 hours. Invasion is quantified as fold area increase in the size of each spheroid over 8 hours. Scale bar = 100 μm. Error bars, SEM. ns, not significant; *p<0.05; ** p<0.01; *** p<0.001.

Furthermore, there was an increased number of large colonies, which is a phenotype typically observed in cells with defects in migration away from growing colonies (Fig. 5E) ^42, 67, 68^. This phenotype is similar to that which we previously observed with Rab27a-KD HT1080 cells ^42^.

To determine whether alterations in exosomes were responsible for the effect of endoglin *in vivo*, we tested whether coinjection of SEVs with the cells could rescue the metastatic defects of endoglin-KD cells. The number of SEVs that were injected with cells was determined using the estimated SEV secretion rate calculated from nanoparticle tracking analysis (NTA) data of 9.28 SEVs per cell per hour for HT1080 control cells. Since the chick CAM experiment was conducted over a period of 96 hours, and each injection contained 100,000 cells, we used 89 million SEVs per injection. Indeed, coinjection of SEVs led to a full rescue of the endoglin-KD phenotype, both increasing the number of metastatic colonies and decreasing the number of large colonies (Fig. 5B-E).

Given the known role of filopodia in directed cell migration ^5^ and the presence of large, potentially nonmotile, colonies formed by endoglin-KD cells in the chick CAM, we hypothesized that endoglin-KD cells would have a defect in cellular migration. To test this possibility, control and endoglin-KD HT1080 cells were allowed to form spheroids and then embedded in 3D type I collagen. Movies were taken to observe and quantitate cell migration away from the spheroids (Supplemental Movie 4). Comparison of the spheroid area increase after 8 h, including migration of individual cells away from the spheroids, revealed that endoglin-KD cells indeed have a defect in 3D cell migration compared to controls. This migratory defect was rescued by re-expression of WT endoglin in the KD cells, confirming that it was not due to an off-target effect of the shRNA (Fig. 5F, Supplemental Movie 4).

### Endoglin controls levels of the filopodia regulator THSD7A in cancer cell SEVs

Endoglin is not expressed in neurons ^69^; therefore, it could not be a universal filopodia regulator. In addition, as a TGFβ coreceptor, it is difficult to imagine how endoglin as an EV cargo could directly induce filopodia formation by recipient cells. However, endoglin interacts with numerous proteins, including TGFβ receptors Alk1 and Alk5, TGFβ family ligand BMP9, and α5β1 integrin, and could potentially alter their sorting into exosomes ^62, 65, 70–72^. Thus, we hypothesized that endoglin could promote filopodia formation by carrying a key filopodia-inducing cargo into exosomes. Surprisingly, Western blot analysis of control and endoglin-KD SEVs did not reveal reductions in several candidate endoglin binding partners in KD SEVs, including TGF-β1, Alk1, or β1 integrin (Fig. S7A). As BMP9 is the only TGFβ family member reported to directly bind to endoglin ^73^, we also tested whether BMP9 could affect filopodia formation. Contrary to expectation, purified BMP9 did not rescue the defect in filopodia formation by endoglin-KD cells and instead decreased filopodia numbers in control shScr cells (Fig. S7B). We also tested whether the addition of TGF-β1 or coating the culture surfaces with fibronectin might rescue the filopodial defect of endoglin-KD cells. While both of those ligands slightly enhanced the number of filopodia in control cells, there was no effect on endoglin-KD cells (Fig. S7C,D).

To determine whether endoglin may regulate the transport of any novel proteins into exosomes, we performed a quantitative proteomic comparison between SEVs purified from control and endoglin-KD B16F1 cells. Equal amounts of protein extracted from the SEVs were subjected to isobaric tagging for relative and absolute quantitation (iTRAQ) and mass spectrometry analysis ^74, 75^. Consistent with our Western blot results (Fig. S7A), the proteomics data showed no significant differences in the expression of any integrins or TGFβ family ligands or receptors (Supplemental Table 2). Analysis of the data further revealed only 2 proteins that were significantly lower in both endoglin KD1 and KD2 samples compared to control samples (Supplemental Table 2), endoglin and the extracellular matrix (ECM) protein thrombospondin type 1 domain containing 7a protein (THSD7A). THSD7A is a transmembrane protein that is known to be expressed in endothelial cells, neurons, and kidney podocytes ^76–79^. Interestingly, although little is known about this protein, a soluble form of THSD7A was reported to promote filopodia formation in endothelial cells ^77^.

Additionally, THSD7A has been shown to be aberrantly expressed in a variety of cancers and autoantibodies against THSD7A are a frequent cause of a paraneoplastic autoimmune kidney disease, secondary membranous nephropathy ^80–82^. Autoantibodies against THSD7A are also a cause of idiopathic (primary) membranous nephropathy ^83^.

To validate our finding that THSD7A levels are downregulated in SEVs from endoglin-KD cancer cells, we performed Western blot analysis of control and endoglin-KD SEVs from B16F1 and HT1080 cells (Fig. 6A,B). Detection of THSD7A in B16F1 SEVs necessitated using native gels and thus endoglin and SEV marker CD63 ran at a significantly higher molecular weight (Fig. 6A). Indeed, THSD7A was present on control SEVs and was reduced on endoglin-KD SEVs purified from either cell line. THSD7A was also present on SEVs isolated from primary cortical neurons (Fig. 6C). To test whether THSD7A regulates filopodia formation in our system, we overexpressed THSD7A tagged with mScarlet fluorescent protein in HT1080 cells. THSD7A-mScarlet localized to the tips of filopodia as well as in extracellular deposits similar to those we have previously observed with exosomes (Fig. 6D ^42, 49^). Notably, expression of THSD7A-mScarlet increased filopodia numbers in HT1080 cells (Fig. 6D, graph). To further test the role of THSD7A in filopodia formation, we knocked down THSD7A in HT1080 cells using 3 different targeting shRNAs (Fig. 6E). Analysis of filopodia numbers revealed a significant reduction in THSD7A-KD cells (Fig. 6E). To determine the role of THSD7A in filopodia regulation by endoglin, we plated control and endoglin-KD cells on coverslips coated with various concentrations of recombinant THSD7A protein. Indeed, recombinant human THSD7A fully rescued the filopodia defects of endoglin-KD for both HT1080 and B16F1 melanoma cells (Fig. 6F).

**Figure 6.**
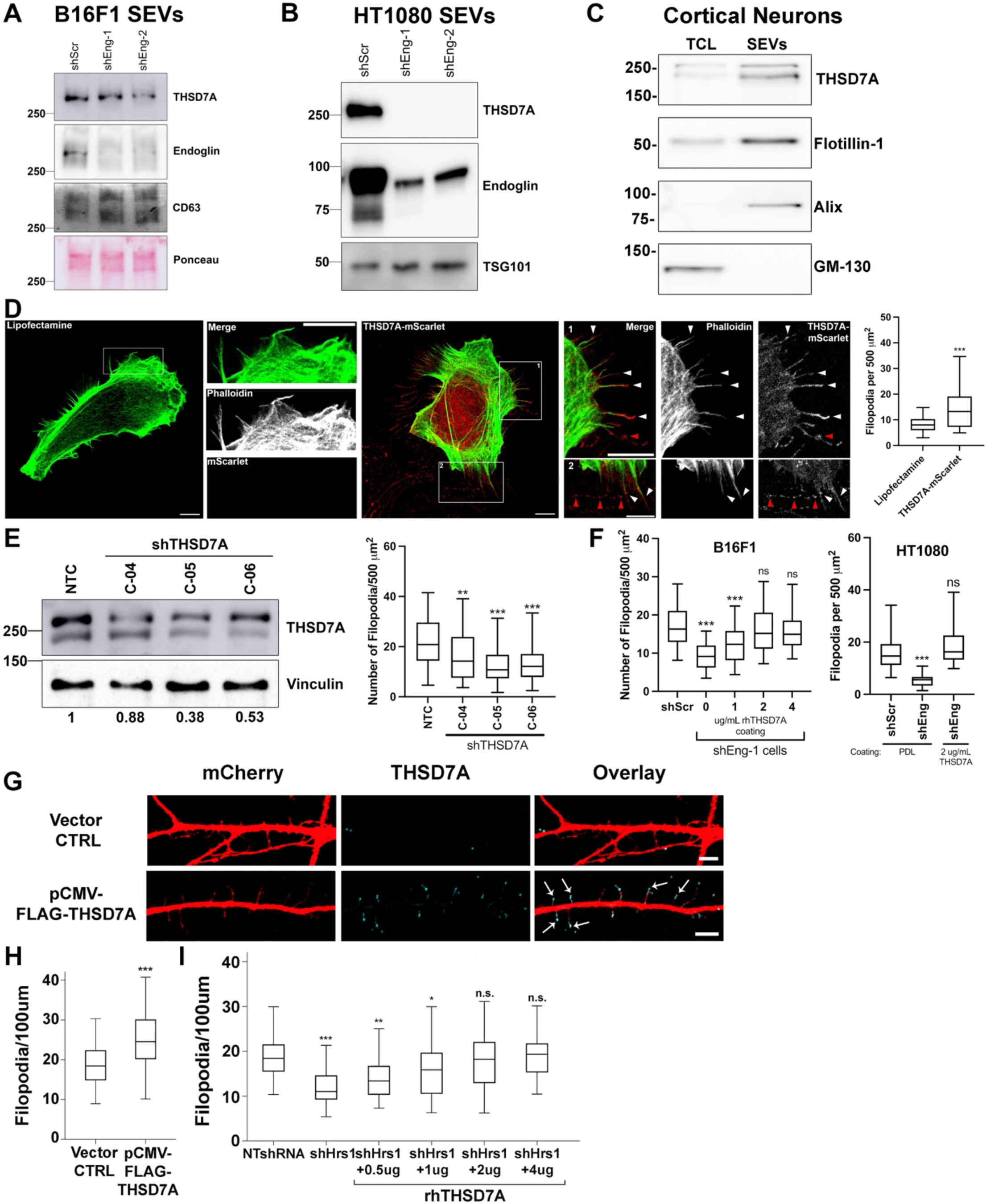
Exosomal endoglin promotes filopodia formation through THSD7A. A. Native gel Western blot of B16F1 SEVs. B. Standard Western blot of HT1080 SEVs. C. Western blot of cortical neuron total cell lysate (TCL) and SEVs. D. Representative images and quantitation of filopodia number in control (lipofectamine) and THSD7A-mScarlet-transfected HT1080 cells. Arrowheads indicate THSD7A at the ends of filopodia (white arrowheads) or in extracellular deposits (red arrowheads). Scale bars in wide field and zoom insets = 10 μm. E. (left) Western blot of control shRNA (NTC) and shTHSD7A (C-04, C05, C-06) - expressing HT1080 cell lines. Vinculin is used as a loading control and numbers below the blot indicate normalized THSD7A levels. (right) Filopodia counts in control and shTHSD7A HT1080 cells. N=3, at least 20 cells per condition per rep. F. THSD7A coated coverslips rescue filopodia defect in shEng B16F1 and HT1080 cells. N=3, at least 20 cells per condition per rep. G. and H. Cortical neurons were transfected with a FLAG-THSD7A expression vector ^77^ or vector control, fixed and stained with an antibody against THSD7A, and imaged by confocal microscopy. G. Representative images. Arrows indicate THSD7A localization to the tips of filopodia. Scale bar = 5 μm. H. Quantification of filopodia in neurons expressing FLAG-THSD7A or control vector. n=42 neurons from three separate experiments. I. Rescue of filopodia numbers in shHrs neurons plated on dishes coated with various concentrations of recombinant human THSD7A, as indicated. Error bars, SEM. ns, not significant; * p<0.05; ** p<0.01; *** p<0.001.

Our typical filopodia analysis is performed after culturing cells for 48 h on coverslips. To determine whether THSD7A can boost filopodia formation in a short time period as would be expected for a direct regulator, we plated cells on coverslips coated with poly-D-Lysine or recombinant THSD7A for 15 min-2 h and found that recombinant THSD7A fully rescued the filopodia defect of endoglin-KD cells at the shortest timepoint of 15 min (Fig. S7E).

Although neurons do not express endoglin ^69^, they do rely on exosomes for filopodia induction (Fig. 3) and they express THSD7A (Fig. 6C, ^78, 84^). To test the role of THSD7A in neuronal filopodia formation, we transiently expressed FLAG-tagged THSD7A in cortical neurons. Immunostaining analysis revealed that neurons expressing FLAG-THSD7A had localization of the THSD7A to filopodia tips and increased numbers of filopodia compared to vector control transfected neurons (Fig. 6G,H). In addition, plating exosome-inhibited Hrs-KD neurons on recombinant THSD7A-coated coverslips fully rescued filopodia formation in a dose-dependent manner (Fig. 6I). These data indicate that THSD7A is an important filopodia-inducing SEV cargo in diverse cell types.

### Endoglin regulates THSD7A trafficking into cancer cell exosomes

Based on the observed decreases in THSD7A expression in SEVs from endoglin-KD cancer cells (Fig. 6A,B) and the role of endoglin as a co-receptor that traffics through endosomes, we hypothesized that endoglin may alter the trafficking of THSD7A to SEVs/exosomes. To test this hypothesis, we first investigated the relative abundance of THSD7A in control and endoglin-KD HT1080 cells and SEVs by Western blot analysis. Indeed, lysates of endoglin-KD cells had increased levels of THSD7A compared to control cells, while endoglin KD SEVs had decreased levels of THSD7A (Fig. 7A-C). These data are consistent with an alteration in THSD7A trafficking in endoglin-KD cells rather than a decrease in the overall protein expression of THSD7A. We performed a rescue experiment, in which endoglin-KD cells were stably transfected with either wild-type endoglin (+WT) or an empty vector control (+vector). Interestingly, we found that the high cellular levels of THSD7A in endoglin-KD cells were decreased with re-expression of WT endoglin (Fig. 7A-C). These changes were paralleled by increased THSD7A in +WT endoglin rescue SEVs. In parallel, expression of WT endoglin in HT1080 endoglin KD cells also rescues filopodia defects (Fig. 7D).

**Figure 7.**
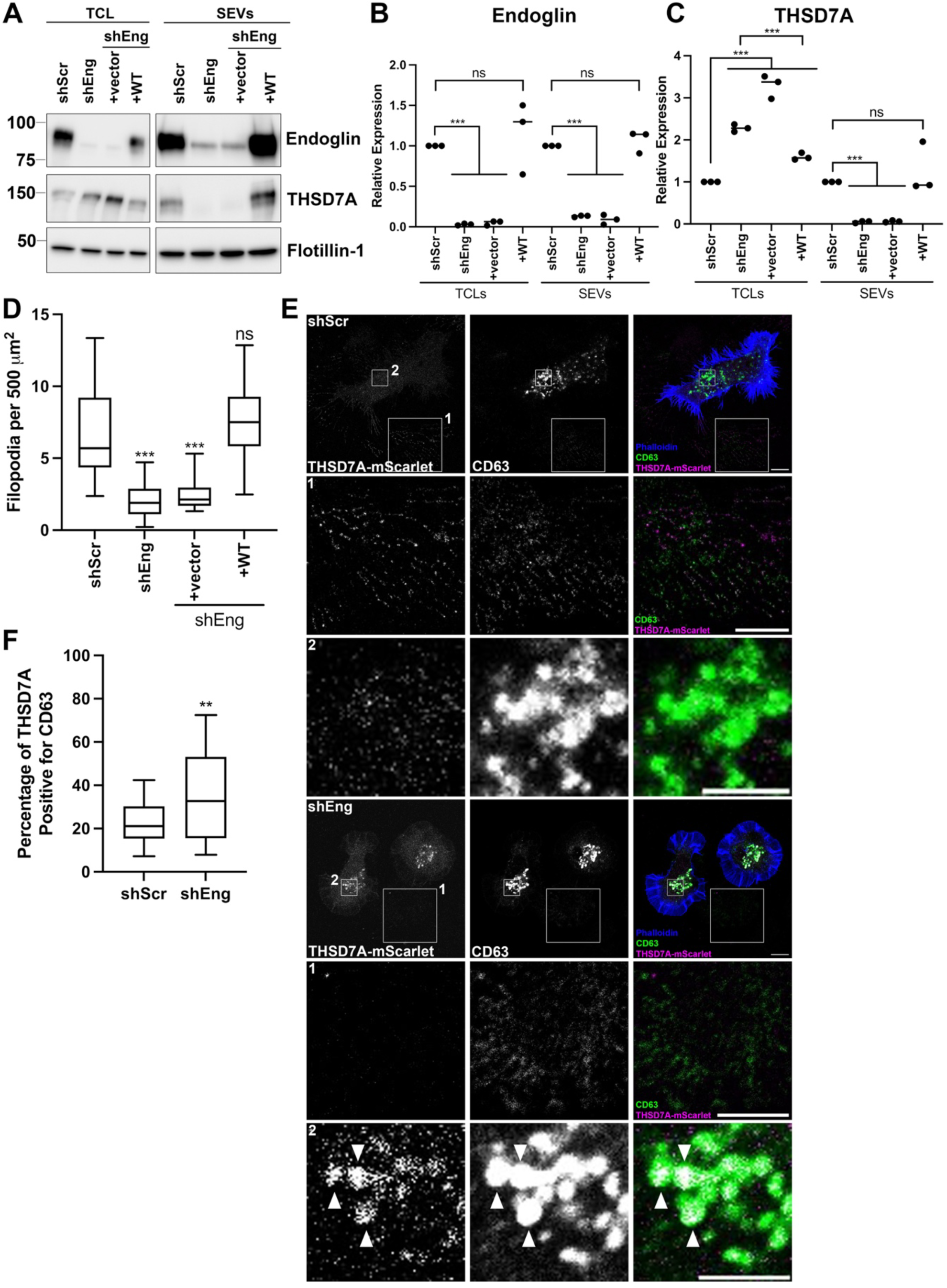
Endoglin controls THSD7A trafficking to exosomes. A. Western blot analysis of total cell lysates (TCL) and SEVs from HT1080 control and shEng cells +/-rescue with WT endoglin or control expression vectors. The figure was made from cropped images of membranes to remove irrelevant lanes. B. Quantification of endoglin expression (normalized to flotillin-1 as a loading control, and relative to shScr control) from triplicate Western blots as in (A). C. Quantification of THSD7A expression (relative to flotillin-1 as a loading control, and relative to shScr control) from triplicate Western blots as in (A). D. Quantification of filopodia in HT1080 control cells and shEng cells rescued with WT endoglin expression. N=3, at least 30 total cells per condition. E. Representative confocal images of THSD7A-mScarlet-expressing control and shEng HT1080 cells immunostained for CD63. Box 1 shows extracellular THSD7A and CD63 deposits. Box 2 shows intracellular CD63-positive MVEs. For both boxes, the zoomed images have been adjusted for brightness and contrast (to equivalent levels for control and shEng cells) for easy visualization. Note that overlap of THSD7A (magenta) and CD63 (green) gives a white signal, pointed out with white arrowheads in shEng merged image in Zoom 2. Scale bar is 10 μm in wider field view and 5 μm in zoom insets. F. Quantification of colocalization of internal CD63 and mScarlet signals in HT1080 cells from nonadjusted images. N=3, ≥ 20 cells per condition per rep. Error bars, SEM. ns, not significant; * p<0.05; ** p<0.01; *** p<0.001.

To determine whether endoglin might affect the trafficking of THSD7A to MVE, we expressed THSD7A-mScarlet in control and endoglin-KD HT1080 cells and co-stained with an anti-CD63 antibody to label MVEs as well as with phalloidin to visualize the cell boundary (Fig. 7E). Consistent with less THSD7A being secreted in EVs, there was much less extracellular THSD7A surrounding endoglin-KD cells compared to control cells (Fig. 7E, Zoom 1). In addition, we observed increased accumulation of THSD7A in CD63-positive MVE in endoglin-KD cells compared to control cells (Fig. 7E, Zoom 2). Thus, the percentage of intracellular THSD7A colocalized with CD63-positive compartments was increased in KD cells compared with control cells (Fig. 7F), suggesting increased accumulation in endolysosomal compartments. These data suggest that endoglin alters the endolysosomal trafficking of THSD7A, leading to decreased THSD7A release in exosomes.

### Cdc42 activity is required for filopodia induction by THSD7A and endoglin

Finally, we sought to identify downstream signaling pathways responsible for modulating filopodia dynamics via exosomal endoglin and THSD7A. To assess any changes in TGFβ signaling, we assayed for downstream Smad phosphorylation and did not detect any increases from treatment with rhTHSD7A coating (Fig. S7F). Because the small GTPase Cdc42 has been shown to regulate filopodia across diverse cell types ^1, 23^, we hypothesized that THSD7A and endoglin might induce filopodia formation via Cdc42 activation. To test this hypothesis, we determined whether filopodia induced by THSD7A could be inhibited with a specific Cdc42 inhibitor or, conversely, whether expressing a constitutively active form of Cdc42 could rescue the filopodia defects of endoglin-KD cells (Fig. 8). The Q61L mutant of Cdc42 (Cdc42-Q61L) is a dominant active, GTPase-defective, GTP-bound form of Cdc42 ^85^. ML141 has been identified as a non-competitive inhibitor of Cdc42 and effectively inhibits Cdc42-Q61L^86^. Consistent with a role for Cdc42 activation in basal filopodia formation by our cells, we found that treatment of control HT1080 cells with the ML141 inhibitor significantly reduces filopodia numbers (Fig. 8). Likewise, the rescue of filopodia numbers that occurs when endoglin-KD cells are seeded on THSD7A-coated coverslips is ablated when combined with ML141 treatment (Fig. 8). In addition, the filopodia defect of endoglin-KD cells is rescued by expression of Cdc42-Q61L. The filopodia induction by Cdc42-Q61L in HT1080 endoglin-KD cells was inhibited by ML141, confirming that ML141 acts to inhibit Cdc42 at the doses and times used (Fig. 8).

**Figure 8.**
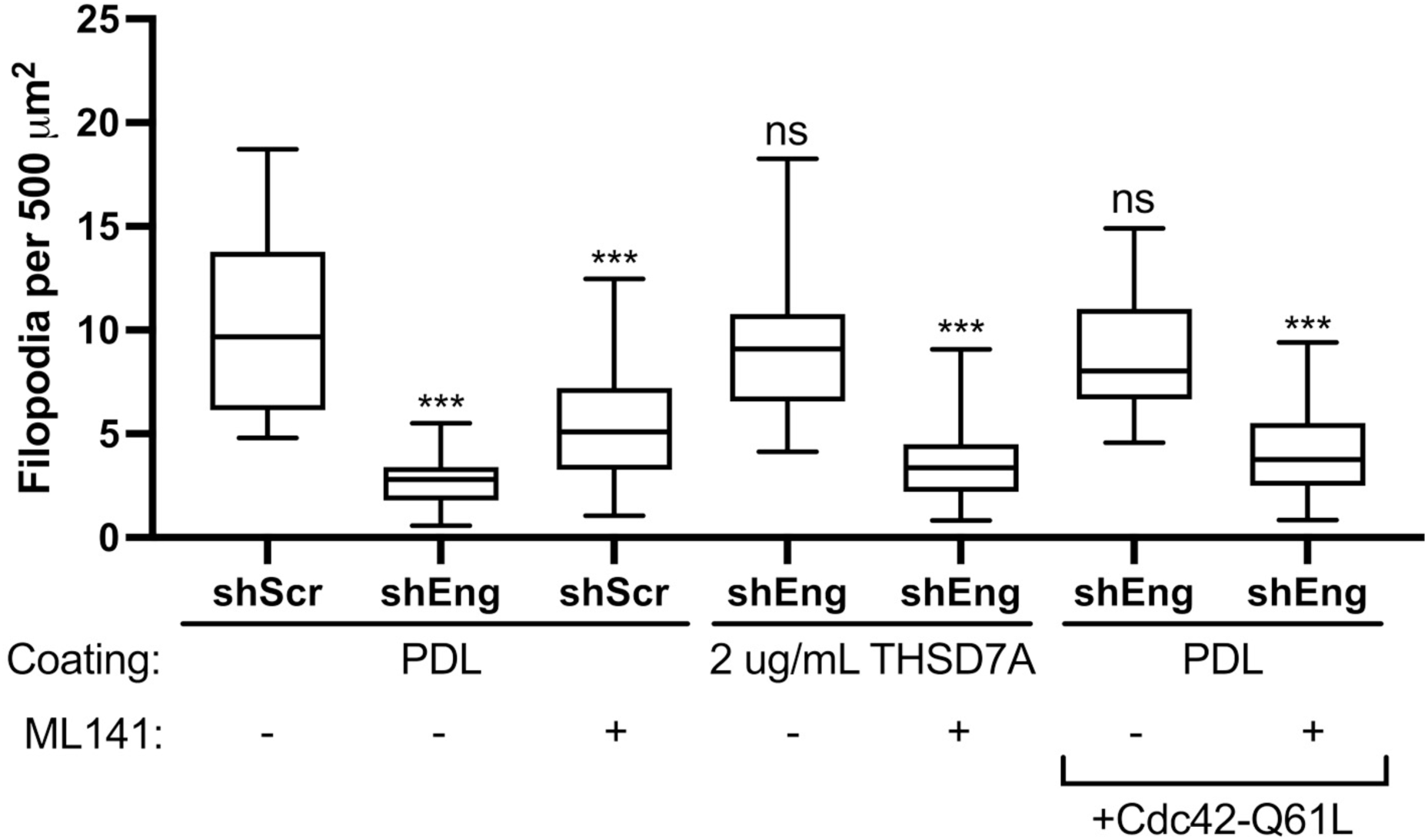
Filopodia formation induced by THSD7A depends on Cdc42 activity. Control and endoglin-KD HT1080 cells were plated on coverslips coated with poly-D-lysine (PDL) or THSD7A. In some cases, cells were treated with the Cdc42 inhibitor ML141 (10 µM) or transfected with the dominant active Cdc42 mutant Q61L, as indicated. N=3, ≥ 20 cells per condition per rep. Error bars, SEM. ns, not significant; * p<0.05; ** p<0.01; *** p<0.001.

## Discussion

Filopodia are adhesive cellular structures that control cell polarization, chemical and physical sensing, and motility. Using a genetic inhibition and rescue approach, we found that autocrine secretion of exosomes controls the number and stability of filopodia in both cancer cells and neurons. The decrease in filopodia in neurons was paralleled by a similar decrease in synapse formation. In cancer cells, we identified endoglin as a key SEV cargo and molecular regulator of this process, controlling filopodia formation, cell migration, and metastasis of cancer cells via SEVs. As endoglin seemed unlikely to directly induce filopodia formation and is not present in neurons, we performed quantitative proteomics of control and endoglin-KD SEVs and identified the transmembrane ECM protein THSD7A as regulated in cancer cell SEVs by endoglin. In neurons, THSD7A was also present on SEVs and regulated filopodia formation. Indeed, purified THSD7A was able to rescue filopodia defects in both endoglin-KD cancer cells and exosome-inhibited neurons. Finally, we found that Cdc42 activity is important for filopodia formation controlled by endoglin and THSD7A. Altogether, we identify THSD7A carried by exosomes as a key controller of filopodia formation in diverse cell types and further identify endoglin as a key regulator of THSD7A secretion in exosomes in cancer cells. These data suggest a new model for filopodia induction via secreted exosomes (Fig. 9).

**Figure 9.**
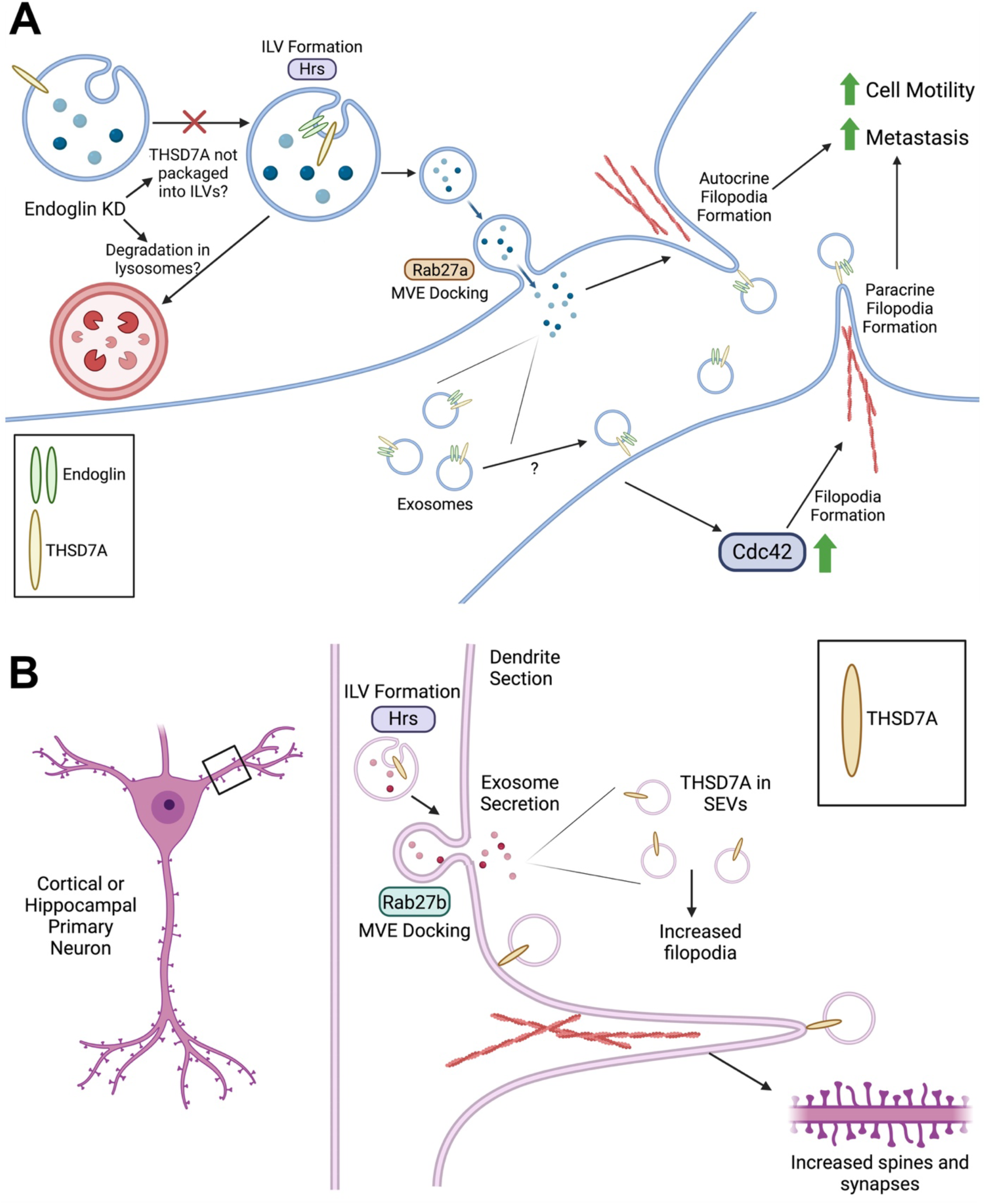
Model of exosome-induced filopodia formation. A. In tumor cells, endoglin and THSD7A are trafficked into intralumenal vesicles (ILV) in multivesicular endosomes (MVEs) for secretion. Inhibiting the exosome biogenesis pathway by blocking Hrs or inhibiting MVE docking by blocking Rab27a subsequently reduces exosome secretion and filopodia formation. SEVs carrying THSD7A can induce filopodia on target cells via Cdc42, leading to increased cell motility and metastasis. When endoglin levels are lowered (such as by KD), THSD7A is retained inside cells in CD63-positive endosomes and its levels in SEVs are greatly decreased. The drop in THSD7A levels in endoglin-KD EVs could be due either to a lack of trafficking into ILVs or, alternatively, enhanced lysosomal degradation of THSD7A-containing MVEs. Given that THSD7A accumulates in CD63-positive endolysosomal compartments in endoglin-KD cells (Fig 7E,F), the latter possibility seems more likely. The cartoon was created using BioRender. B. In primary neurons, exosome biogenesis is controlled by the formation of ILVs by Hrs and MVE docking is controlled by Rab27b. Knockdown of either of these proteins results in reduced formation of filopodia, dendritic spines, and synapses in both cortical and hippocampal neurons. Similar to cancer cells, THSD7A is carried in neuronal SEVs and induces filopodia. The cartoon was created using BioRender.

Exosomes have been shown to regulate cell migration and invasion in a variety of contexts ^40–44, 49^. In order for cells to migrate, they must form adhesive and sensing cytoskeletal structures, including lamellipodia, focal complexes, invadopodia, and filopodia. Our finding that exosomes promote filopodia formation adds to our previous findings that exosome secretion promotes the formation of nascent adhesions and invadopodia ^42, 48^ and likely stabilizes lamellipodia ^87–89^. While some of these activities are likely related, the filopodia activity appears to be distinct. Indeed, while we previously found that fibronectin was a critical exosome cargo driving nascent adhesion formation, lamellipodia stabilization, and cell speed in HT1080 fibrosarcoma cells ^42, 88, 89^, fibronectin was unable to rescue the filopodia defect of endoglin-KD HT1080 cells in this study (Fig. S7D). Instead, THSD7A was identified as a distinct exosome cargo that specifically promotes *de novo* filopodia formation in HT1080 and other cell types and rescued defects of exosome-inhibited cells in filopodia formation. Thus, multiple exosome cargoes contribute to diverse aspects of cytoskeletal reorganization and cell migration.

While no study has previously shown that EVs can induce filopodia formation, filopodia have previously been shown to interact with EVs in diverse ways. EVs were shown to bind and “surf” on filopodia before entering cells ^90^. EVs have also been shown to bud from the surface of filopodia and other similar protrusions ^36, 91–94^. These various interactions influence cells in a variety of ways, including the delivery and release of EV cargoes and the promotion of cellular migration and invasion.

Filopodia are important for directional sensing, directional migration, and cell-cell contact formation in a variety of cell types ^1, 4, 5^. In cancer, filopodia formation is associated with aggressive behavior and cancer metastasis. Indeed, we found that endoglin-KD cancer cells had defects in not only filopodia formation but also in 3D migration and metastasis to chick embryo chorioallantoic membranes. In neurons, inhibition of exosome secretion reduced both filopodia and synapse formation. Thus, in two very different cell types, exosomes regulate these fundamental, filopodia-dependent behaviors. Future studies could further test the importance of exosomes and exosome-associated-THSD7A in controlling filopodia in additional cell types, including endothelial cells that use filopodia during cell-cell contact formation and angiogenesis ^95^. It would also be interesting to see whether THSD7A carried by EVs regulates kidney podocyte foot process formation downstream of filopodia and whether it is soluble proteolyzed THSD7A or THSD7A-carrying EVs released into the circulation by cancer cells that leads to autoimmune secondary membranous nephropathy ^76, 82^.

In our first proteomics dataset identifying SEV-enriched cargoes, we identified endoglin as a candidate molecule to regulate filopodia formation. While we originally hypothesized that endoglin was regulating filopodia formation through one of its known binding partners, our tests of candidates did not yield any positive results. Instead, through our proteomic comparison of control and endoglin-KD SEVs, we identified THSD7A as a possible filopodia-inducing EV cargo regulated by endoglin. Indeed, recombinant purified THSD7A rapidly induced filopodia formation, rescuing filopodia defects of endoglin-KD cells within 15 min (Fig. S7E). As neurons do not express endoglin ^69^ but do express THSD7A, we expect that sorting of THSD7A to exosomes is controlled by additional molecules. In addition, we found for the first time that Cdc42 activity was required for THSD7A to promote filopodia formation. However, the intermediary proteins that link THSD7A to Cdc42 remain unknown. Future studies to identify THSD7A-binding partners will be important to understand both the mechanism by which THSD7A selectively induces filopodia formation via Cdc42 and how its trafficking is regulated in neurons and other cell types.

Endoglin is highly expressed in endothelial cells and regulates angiogenesis ^63^. Indeed, mutations in endoglin are a frequent cause of hereditary hemorrhagic telangiectasia – a disease in which abnormal vascular structures are formed in the skin, mucous membranes and some organs ^96^. Endoglin expression is also upregulated in several cancers ^97^, including melanoma ^98^, ovarian cancer ^99^, breast cancer ^100^, and gastric cancer ^101^. In breast cancer, these elevated endoglin levels are correlated with reduced survival and invasive phenotype ^100, 102^. There is also blood vessel enrichment of endoglin in ovarian tumors ^103^ and head and neck squamous cell carcinoma ^104^. In fact, anti-endoglin antibodies are being evaluated as potential anti-cancer therapy and inhibit angiogenesis and metastasis in pre-clinical models ^105–108^. Given our new findings that endoglin is a key regulator of filopodia via regulating THSD7A trafficking to SEVs, an important future direction could be to test the contribution of THSD7A to endoglin-dependent phenotypes, including angiogenesis and cancer metastasis.

In summary, we found that exosomes promote filopodia formation in diverse cell types by carrying the transmembrane ECM molecule THSD7A. This appears to be a major mechanism for filopodia formation with broad importance for a variety of cellular processes, including cancer cell migration and neuronal synapse formation.

## Limitations of the Study

The main findings of this study are 1) exosomes induce filopodia formation in both cancer cells and neurons and synapse formation in primary neurons; 2) endoglin and THSD7A are exosomal cargoes that promote filopodia formation; 3) endoglin regulates the levels of THSD7A in cancer cell SEVs and promotes cancer metastasis; 4) THSD7A is expressed by both cancer cells and neurons and directly induces filopodia formation, including in cells that cannot secrete exosomes due to genetic engineering. As a first step to identify the mechanism by which THSD7A induces filopodia formation, we tested whether the candidate molecule Cdc42 is required downstream of THSD7A. We used the inhibitor ML141 and a dominant active Cdc42 molecule to show respectively that induction of filopodia by THSD7A is inhibited with ML141 and that activation of Cdc42 can overcome the filopodia defects of endoglin-KD. One limitation of the study is that we did not use a biochemical method to show that exosomes can activate Cdc42 in cells. Activation of Cdc42 is typically quantified by assessing the ratio of the GDP-bound inactive form to the GTP-bound active form via methods such as bead pulldown assays and Western blots. We were unable to detect active Cdc42 in our cancer cell lysates, suggesting the need to greatly scale up and further optimize in order to achieve the required signal for this assay. Thus, while our data show promising results to support the role of Cdc42 in this process, further experimentation using orthogonal methods such as Cdc42 activity assays and testing Cdc42 knockdown would strengthen this finding and be an important future direction. Another future direction could be to test the role of additional proteins known to be involved in filopodia formation, such as actin-binding proteins like mDia or ENA/VASP proteins, or BAR domain proteins. A second limitation is that we have not worked out the full mechanism by which endoglin regulates THSD7A secretion in exosomes. In order to do so, we would need to determine definitively how THSD7A trafficking is altered by endoglin, i.e. alterations in secretion vs. formation of THSD7A-containing EVs (Fig 9A). More in-depth assessments of the cellular distribution of THSD7A and endoglin, along with trafficking assays, would aid this endeavor.

## Materials and Methods

### Cancer Cell Methods

#### Cell Culture

B16F1 cells were maintained in DMEM supplemented with 10% FBS. HT1080 cells were maintained in DMEM supplemented with 10% bovine growth serum (BGS). Cells were maintained at 37 degrees in 5% CO2. Transient transfections were done using Lipofectamine 2000 (Invitrogen #11668-027).

#### Plasmids and Reagents

Stable knockdown of Hrs in B16F1 cells was achieved by using Dharmacon p-Blockit system with target sequences: 5’-GGAACGAACCCAAGTACAAGG-3’ (shHrs-1) and 5’-GCATGAAGACGAACCACATGC-3’ (shHrs-2) and stably selected using blasticidin. Stable knockdown of Rab27a in B16F1 cells was achieved by using Dharmacon p-Blockit system with target sequences: 5’-GTGCGATCAAATGGTCATGCC-3’ (shRab27a-1) and 5’-CGTTCTTCAGAGATGCTATGC-3’ (shRab27a-2) and stably selected using blasticidin. Control lines were simultaneously selected using the pBlockit-shLacZ plasmid. Stable knockdown of endoglin in B16F1 cells was achieved using Dharmacon TRC lentiviral shRNA (pLKO.1): 3’UTR target shEng-1 (TRCN0000094354): 5’-TTAGGCTTCTAAGCAGCATGG-3’, ORF target shEng-2 (TRCN0000094355): 5’-TATAGATGACAAACAGCAGGG-3’. B16F1 shEng cells were stably selected using puromycin. The control line was simultaneously selected using the shScr control plasmid. Transient endoglin knockdown in B16F1 cells was obtained with ON-TARGETplus siRNA SMARTpool (L-045109-00-0005; GE Dharmacon) using Lipofectamine RNAiMAX (Life Technologies). As a control for the knockdown, a nontargeting control pool was used (D-001810-01-05; GE Dharmacon). Stable knockdown of endoglin in HT1080 (mCitrine expressing) cells was achieved using Dharmacon TRC lentiviral shRNA (pLKO.1): 3’UTR shEng-1 (TRCN0000083138): 5’-ATCCAGGTTCAAATGACAGGG-3’, ORF shEng-2 (TRCN0000083141): 5’-ATCATACTTGCTGACACCTGC-3’, CDS shEng (TRCN0000083139): 5’-TAGTGGTATATGTCACCTCGC-3’. Cells were selected with puromycin and maintained in blasticidin and puromycin to retain mCitrine expression and endoglin knockdown. The control line was simultaneously selected using the shScr control plasmid. Knockdown of THSD7A in HT1080 cells was achieved using Dharmacon SMARTvector Lentiviral shRNAs Cat# V3SH11240-226004853, V3SH11240-226644723, and V3SH11240-227287134. A non-targeting control (NTC) shRNA was used as a control. pCMV-Tag4-THSD7A-FLAG plasmid and vector control pCMV-Tag4 were generously provided by the Chuang laboratory^77^. THSD7A-mScarlet plasmid was created by cloning mScarlet region from pCytERM_mScarlet_N1 (Addgene #85066) and inserting it at the C terminal end of THSD7A in the pCMV-THSD7A plasmid obtained from Chuang lab. For mechanistic studies, HT1080 shEng cells were transfected with empty control vector pLenti6/V5-DEST or wild type endoglin pLenti6-WT-Eng-V5. The blasticidin resistance gene in pLenti6/V5-DEST was replaced with neomycin resistance gene. The WT endoglin plasmid had multiple silent mutations inserted to confer resistance to the endoglin shRNA. Stable lines were created using selection with G418 sulfate. pRK5-myc-Cdc42-Q61L was purchased from Addgene (Plasmid #12974). HT1080 cells stably expressing mCitrine fluorescent protein were obtained from Andries Zijlstra’s laboratory. HT1080 cells stably expressing the dual tagged CD63 live imaging reporter pHluorin_M153R-CD63-mScarlet were previously described ^109^.

#### EV Isolation and Characterization

48-hour conditioned Opti-MEM (a serum-free but growth factor-containing medium) was collected for isolation of EVs from cancer cells. A 5-minute 300xg spin and 25-minute 2000xg spin were performed to remove dead cells and large particulate matter. The supernatant was then spun at 10,000xg for 30 minutes to pellet the LEVs (Ti45 rotor, Beckman Coulter). This LEV fraction was later washed with PBS spins and used for Western blotting and rescue experiments. To isolate SEVs from cancer cells, we used a cushion density gradient ultracentrifugation method, which reduces vesicle and protein aggregation and leads to a highly purified preparation ^54^. For iodixanol cushion and gradient, OptiPrep Density Gradient Medium was purchased from Sigma (D1556). The supernatant from the 10,000xg spin was layered on top of a 2 mL 60% iodixanol cushion and spun for 4 hours at 100,000xg (SW32 Ti rotor, Beckman Coulter). The bottom 3 mL of the cushion was obtained and layered at the bottom of a discontinuous gradient, followed by three 3 mL layers of 20%, 10%, and 5% iodixanol diluted with 0.25M sucrose/10mM Tris, pH 7.5. The discontinuous gradient was spun for 18 hours at 100,000xg and collected into twelve 1 mL fractions (SW40 Ti rotor, Beckman Coulter). SEVs are typically located in fractions 6 and 7, and these fractions are then washed by resuspending with PBS and repelleting. After final resuspension with PBS, SEV purification is checked using nanoparticle tracking for particle number and size, Western blot validation of EV proteins, and electron microscopy for morphology ^55–57^. Large volumes of conditioned media were concentrated prior to the cushion centrifugation step using Vivacell 70 ultrafiltration units with 100,000 MWCO. The number of vesicles added back in rescue experiments was determined by calculating the physiological SEV secretion rate for each cell line. The total vesicle number was counted using nanoparticle tracking analysis (Particle Metrix ZetaView or NanoSight), and the vesicle per cell per hour secretion rate was calculated. This rate was used to determine a range of physiologically relevant vesicle numbers to treat cells.

#### Electron Microscopy

Freshly isolated SEVs or LEVs suspended in PBS were used for TEM imaging. Samples were incubated on glow discharged 300 mesh carbon coated grids for 30 seconds followed by fixation in 1% glutaraldehyde for 1 minute. Samples were washed twice and negative stained in 2% uranyl acetate. TEM was performed on a Tecnai T12 operating at 100 keV using an AMT nanosprint5 CMOS camera.

#### Immunofluorescence and Analysis

Cells were seeded onto 100 ug/mL PDL-coated glass coverslips. For steady-state filopodia quantification, cells were fixed with 4% paraformaldehyde, permeabilized with 0.1% Triton X-100 or 0.1% saponin, and stained with Alexa Fluor conjugated phalloidin (rhodamine-phalloidin or phalloidin-Alexa Fluor 488) 24 hours after seeding. Co-staining was done using mouse anti-CD63 (Abcam ab8219) and goat anti-mouse Alexa Fluor 546. For EV rescue experiments, SEVs or LEVs are added to live cells 24 hours post-seeding and cells are fixed and stained with conjugated phalloidin 18 hours post-EV treatment. For BMP-9 treatment, human recombinant BMP-9 (R&D Systems Cat# 3209-BP) was reconstituted in sterile 4mM HCl + 0.1% BSA, diluted in OptiMEM and added to cells for 1 hour at 37 degrees C, then cells are fixed and stained with conjugated phalloidin. For TGF-β1 treatment, desired concentration of recombinant human TGF-β1 (R&D Systems Cat #240-B-002; reconstituted in sterile 4 mM HCl + 0.1% BSA) was diluted in OptiMEM, added to cells at time of seeding, and cells were fixed and stained 48 hours post-seeding. For FN coating experiments, glass coverslips were coated with indicated concentrations of FN (Cultrex Human Fibronectin Cat# 3420-001-01, diluted in sterile PBS) overnight and cells were seeded onto coverslips. Cells were fixed and stained 24 hours post-seeding. For THSD7A rescue experiments, coverslips were first coated with 100 ug/mL PDL overnight, then washed, then coated with indicated concentrations of recombinant human THSD7A overnight (diluted in sterile PBS) (R&D Systems Cat# 9524-TH). Cells were seeded onto THSD7A-coated coverslips and fixed and stained 24 hours post-seeding. For THSD7A-mScarlet expression experiments, THSD7A-mScarlet was transiently transfected into HT1080 cells and cells that were visibly expressing the fusion protein were imaged using a Nikon A1R HD25 confocal microscope with a Plan Apo λ 60x/1.4 oil objective. Cells were seeded onto PDL-coated glass coverslips overnight then fixed and stained for phalloidin and CD63. Cell edges were manually drawn and only the internal THSD7A-mScarlet and CD63 signals were quantified. JACoP Fiji plugin was used to calculate M1 value from manually threshholded channels. All fixed cells were imaged using either a Nikon Eclipse TE2000-E epifluorescence microscope and Metamorph software (Molecular Devices) or Nikon A1R confocal microscope and NIS-Elements Software. Cell area was calculated by tracing cell outlines using ImageJ software. Filopodia were counted using Filoquant ImageJ plugin ^16^ with manually adjustment for any plugin errors.

#### Live Cell Imaging

<u>MVE fusion movies:</u> HT1080 cells stably expressing pHluorin_M153R-CD63-mScarlet were cultured on PDL-coated glass-bottomed MatTek dishes. Cells were imaged every 10 seconds for 20 minutes on a Nikon A1R confocal microscope using a Plan Apo λ 60x/1.4 oil objective and a humidified 37-degree C chamber with 5% CO_2_. MVE fusion events are identified as MVEs expressing mScarlet-CD63 approaching the cell edge followed by a burst of green fluorescence, suggesting exposure to extracellular neutral pH and MVE fusion. Filopodia that arose at the region of MVE fusion soon after were counted in the quantification. CD63 green burst and filopodia formation were noted based on the time frame, and the timing between these events was quantified and graphed.

#### Filopodia dynamics

B16F1 control (shLacZ) and Hrs-KD cell lines were transiently transfected with tdTomato-fTractin 2 days prior to seeding onto PDL-coated glass-bottomed MatTek dishes. Prior to imaging, cell media was switched to Leibovitz’s L-15 (Gibco) + 10% FBS. Cells were imaged on Nikon Eclipse TE2000-E epifluorescence microscope using Apo 60x oil objective and warmed 37 degree C chamber and an image was captured every 30 seconds for 15 minutes with MetaMorph software (Molecular Devices). B16F1 control (shScr) and Eng-KD cell lines were transiently transfected with tdTomato-fTractin 2 days prior to seeding onto PDL-coated glass-bottom MatTek plates. Cells were imaged on Nikon A1R confocal microscope using Plan Apo 11 60x/1.4 oil objective and a humidified 37 degree C chamber with 5% CO_2_. An image was captured every 30 seconds for 15 minutes with NIS-Elements software. Only filopodia that were formed and completely retracted during the 15 minute time period were quantified as de novo and lifetime was defined as the total amount of time the filopodia persisted from formation to complete retraction.

### Proteomics Analyses

#### Identification of individual bands isolated from colloidal Coomassie blue-stained gel

Purified LEVs and SEVs isolated from B16F1 cells were run on a standard SDS-PAGE gel using fresh buffers and after washing the gel apparatus and cleaning it with 70% ethanol. After staining and destaining with the BioRad Coomassie Brilliant Blue R-250 Staining Solutions Kit (#1610435), four bands that were apparent in the SEV sample but missing in the LEV sample were identified, cut from the gel, and submitted for proteomics analysis using trypsin as a digestion enzyme. MS/MS samples were analyzed using Sequest (Thermo Fisher Scientific, San Jose, CA, USA; version 27, rev. 12) and X! Tandem (The GPM, thegpm.org; version CYCLONE (2010.12.01.1)). Sequest was set up to search the uniprot-mouse-reference-canonical_20121112_rev database (unknown version, 86222 entries), assuming the digestion enzyme trypsin. X! Tandem was set up to search the uniprot-mouse-reference-canonical_20121112_rev database (unknown version, 86222 entries), also assuming trypsin. Sequest was searched with a fragment ion mass tolerance of 0.00 Da and a parent ion tolerance of 2.5 Da. X! Tandem was searched with a fragment ion mass tolerance of 0.50 Da and a parent ion tolerance of 2.5 Da. Glu->pyro->Glu of the n-terminus, ammonia-loss of the n-terminus, gln->pyro->Glu of the N-terminus, oxidation of methionine and carbamidomethyl of cysteine were specified in X! Tandem as variable modifications. Oxidation of methionine and carbamidomethyl of cysteine were specified in Sequest as variable modifications. Criteria for protein identification: Scaffold (version Scafford_4.8.8, Proteome Software Inc., Portland, OR) was used to validate MS/MS based peptide and protein identifications. Peptide identifications were accepted if they could be established at greater than 50.0% probability to achieve an FDR less than 5.0% by the Peptide Prophet algorithm (Keller, A. et al. Anal. Chem. 2002;74(20):5483-92). Protein identifications were accepted if they could be established at greater than 81.0% probability to achieve an FDR less than 5.0% and contained at least 2 identified peptides. Protein probabilities were assigned by the Protein Prophet algorithm (Nesvizhskii, Al et al. Anal. Chem. 2003;75(17):4646-58). Proteins that contained similar peptides and could not be differentiated based on MS/MS Analysis alone were grouped to satisfy the principles of parsimony. The full proteomics results are shown in Supplemental Table 1.

#### iTRAQ Analysis of SEVs

For iTRAQ proteomics analysis, isolated exosomes from multiple cushion-density gradient preparations were pooled together for each cell type. Each individual preparation was tested for purity using Zetaview nanoparticle tracking and Western blot for typical exosomal markers. Pooled SEVs were resuspended in PBS and mixed 1:1 with 2x “exosome lysis buffer” (200 mM TEAB, 600 mM NaCl, 2% NP-40, and 1% sodium deoxycholate). After mixing, samples were then sonicated using a Bioruptor in cold ice water (30 seconds on/30 seconds off for 15 minutes). After sonication, samples were spun down to pellet insoluble proteins, and the supernatant was submitted and run by the Vanderbilt University MSRC Proteomics Core Laboratory. MS/MS spectra were searched against a mouse subset of the UniProt KB protein database, and autovalidation procedures in Spectrum Mill were used to filter the data to <1% false discovery rates at the protein and peptide level. The median log2 iTRAQ protein ratios were calculated over all peptides identified for each protein, and frequency distributions were generated in GraphPad Prism. Log2 ratios typically follow a normal distribution and were fit using least squares regression. The mean and standard deviation values derived from the Gaussian fit were used to calculate p-values using Z-score statistics. A given iTRAQ protein ratio, with the calculated mean and standard deviation of the fitted dataset, is transformed to a standard normal variable (z = (x-μ)/σ). Since the properties of the standard normal curve are known, area under the curve for a particular value can be calculated, providing a p-value for each measured protein ratio. Calculated p-values were subsequently corrected for multiple comparisons using the Benjamini-Hochberg method. The full proteomics results are shown in Supplemental Table 2.

#### Western Blot Analysis

Samples for Western blotting were run on a reducing SDS-PAGE gel and transferred to a nitrocellulose membrane (unless otherwise noted). Cell lysate samples were collected by scraping cells directly from a tissue culture dish using RIPA cell lysis buffer (50 mM Tris pH 7.6, 150 mM NaCl, 1% NP-40, 1% SDS, 0.5% sodium deoxycholate) with 1 mM Phenylmethylsulfonyl fluoride (PMSF) (Research Products International Cat# P20270) and cOmplete^TM^ Protease Inhibitor Cocktail (Roche Cat# 04693116001, used as directed by manufacturer). EV samples were lysed by mixing directly with Laemmli sample buffer containing SDS, DTT, and BME. Samples were loaded at equal protein amounts; protein amount was quantified from samples by using a BCA assay or micro BCA assay (Pierce^TM^ BCA Protein Assay Kit Thermo Scientific Cat# 23225, Micro BCA^TM^ Protein Assay Kit Thermo Scientific Cat# 23235). Antibodies (for cancer cells): rabbit anti-Endoglin (mouse specific) (#3290, Cell Signaling), rabbit anti-Endoglin (human specific) (#4335, Cell Signaling), rabbit anti-CD63 (ab68418, Abcam), rabbit anti-beta actin (#4970, Cell Signaling), rabbit anti-TSG101 (ab30871, Abcam), rabbit anti-flotillin-1 (#3253, Cell Signaling), rabbit anti-TGFbeta1 (ab92486, Abcam), mouse anti-HSP70 (sc-24, Santa Cruz), mouse anti-CD29 (beta1-integrin; #610467, BD Biosciences), rabbit anti-ALK1/ACVRL1 (Abcepta AP7807a), rabbit anti-Rab27a (#69295, Cell Signaling), rabbit anti-Hrs (M-79) (sc-30221, Santa Cruz), rabbit anti-GM130 (ThermoFisher MA5-35107), rabbit anti-THSD7A (HPA000923, Sigma), mouse anti-vinculin (Sigma-Aldrich V9131).

To detect THSD7A in HT1080 TCLs and SEVs, reducing Western blot conditions and rabbit anti-THSD7A (Sigma HPA000923) were used. To detect THSD7A in B16F1 SEV samples, SEVs were solubilized in NP-40 lysis buffer (50 mM Tris pH 8.0, 150 mM NaCl, 1.0% NP-40, 1mM PMSF mixed 1:1 with SEVs resuspended in PBS) in non-reducing conditions and ran on native gels using non-SDS running buffer. After transfer, the membrane was probed with rabbit anti-THSD7A IgG generously provided by Nicola M. Tomas, Rolf A.K. Stahl, and Friedrich Koch-Nolte at University Medical Center Hamburg-Eppendorf in Hamburg, Germany^110^.

For detecting Smad phosphorylation, TGF-β1 was used as a positive control and added to indicated conditions for 1 hr at 10 ng/mL. TGF-β1 was reconstituted in sterile 4 mM HCl + 0.1% BSA according to manufacturer’s instructions (R&D Systems Cat #240-B-002). Indicated conditions were treated with the TGF-β1 inhibitor SB 431542 (Sigma-Aldrich Cat# S4317; solubilized in DMSO) for 5 minutes at 10 μM prior to TGF-β1 treatment. Cells were seeded on 100 μg/mL PDL +/-2 μg/mL rhTHSD7A. At experiment endpoint, media was aspirated and adherent cells were lysed with RIPA lysis buffer (50mM Tris pH 7.6, 150mM NaCl, 1% NP-40, 1% SDS, 0.5% sodium deoxycholate) containing 1mM phenylmethylsulfonyl fluoride (PMSF) (Research Products International (RPI) Cat # P202705) and PhosSTOP^TM^ (Roche Cat # 4906845001). Lysates were passed through a 27-gauge needle three times prior to boiling and preparing for Western blot. pSmad1/5/9 (Cell Signaling #13820) and Smad1 (Cell Signaling #6944) were probed on one membrane and pSmad2 (Cell Signaling #3108) and Smad2 (Cell Signaling #5339) were probed on a separate membrane. Membranes were blocked with 5% BSA in TBS/0.1% Tween-20 overnight, probed with indicated phospho-antibody and then stripped and reprobed with indicated total Smad antibody.

#### Avian Embryo Model of Metastasis

Avian embryo experimental metastasis model protocol was based on previously published methods ^66^. Live fertilized chicken eggs were incubated for 11 days prior to injections. HT1080 control and KD cells expressing mCitrine were suspended in PBS at a concentration of 1×10^6^ cells/mL and 100,000 cells (100 μL) were injected into the allantoic vein in the direction of bloodflow. Eggs were returned to the incubator for 4 days. After 4 days post-injection, chicks were sacrificed, and three circular areas of CAM were harvested from inside the shell 4 days post-injection. The membrane was then peeled away from the eggshell and placed between a glass slide and coverslip for imaging. 25-30 fields of view were captured for each egg harvested and 4-7 eggs were sacrificed for each condition. Preliminary low-power wide-field images were obtained using a Zeiss Lumar V12 fluorescence stereomicroscope with 10X magnification. For higher power wide-field images, fluorescent cell extravasation and metastasis was observed using X10/0.40 UPlanSApo objective lens with a 10X ocular (100X total) on a Olympus BX-61 microscope equipped with a digital camera controlled with Volocity image acquisition software. For SEV rescue experiments, control SEVs were isolated using the cushion-density gradient method and were premixed with cells immediately prior to injection into chick embryo intravascularly. The number of SEVs per injection was calculated using the SEV secretion rate for the HT1080 shScr cell line according to data collected with the Zetaview nanoparticle tracking instrument (89×10^6^ SEVs per 100,000 cells). Images were analyzed in ImageJ by manually thresholding each image to include visible cells and avoid capturing background signals. The colony number and colony area of each thresholded image were quantified.

#### Spheroid Collagen Invasion Model

1.5 × 10^4^ HT1080 cells were cultured in an EZSPHERE micro-fabricated 96-well plate (AGC Techno Glass, Shizuoka, Japan). After 20 hrs, the spheroids from each well were collected into a 15-ml tube by adding 5 ml of DMEM medium and centrifuging at 300 g for 1 min. For collagen gel culture, a cold 3 mg/ml type 1A collagen solution (Nitta gelatin, Osaka, Japan) was neutralized by adding an 8:1:1 ratio of collagen : 10 x DMEM/F12 : reconstitution buffer (200 mM HEPES, 50 mM NaOH, and 260 mM NaHCO3, then diluted in cold PBS to achieve a final concentration of 1.2 mg/ml. 10 μl of the neutralized collagen/medium solution was coated onto a 35-mm glass bottomed culture dish and solidified at 37 °C for 30 min. The collected spheroids were suspended in 100 μl of the ice-cold collagen/medium solution, domed onto the lower collagen gel, and solidified at 37 °C for 30 min. 2 ml of the complete medium was gently added to the dish, and Time-Lapse images were captured using a Keyence digital fluorescence microscope BZ-9000 using the 10X phase contrast objective lens.

#### Cdc42 Inhibition Assay

Cells were transiently transfected with constitutively active Cdc42-Q61L 24 hours prior to seeding on 100 µg/mL PDL coated coverslips with or without 2 ug/mL rhTHSD7A. 24 hours after seeding, cells were treated with either vehicle (DMSO) or 10 µM ML141 (Millipore Cat# 217708) diluted in OptiMEM media for 60 minutes. After treatment, cells were fixed with 4% paraformaldehyde and stained with phalloidin to visualize filopodia and Hoechst to visualize nuclei.

### Neuron Methods

#### Reagents and Constructs

mCherry cDNA was a generous gift from Roger Tsien (University of California, San Diego, CA). GFP and mCherry were cloned into pTαS2 vector, kind gift from Freda Miller, for expression in neurons. SV2 monoclonal antibodies were obtained from the Developmental Studies Hybridoma Bank (University of Iowa, Iowa city, IA). Alexa Fluor 488 Anti-Rabbit and Alexa Fluor 647 Anti-Mouse were from Molecular probes. For neuronal cultures, B27 media was prepared by adding 2% B27 supplement and L-glutamine to neurobasal media.

#### Primary Cultures of Neurons

Rat hippocampal and cortical neurons were isolated from day 19 embryos and plated on 1 mg/ml Poly-L-Lysine (PLL) or 50ug/ml Poly-D-Lysine coated glass coverslips. Low-density cultures were prepared as described previously. In brief, the hippocampus and cortex was removed from dissected brains of day 19 rat embryos and incubated in 0.05% trypsin in HBSS for 10 mins at 37^0^C. Neurons were washed with HBSS, homogenized by gentle mixing and plated on PLL or PDL coated coverslips. After 3-4 hours, neuron coverslips were transferred to 60mm dishes containing primary astroglia for co-culture to promote neuronal health ^111^.

#### SEV Isolation and Characterization

For neuronal SEVs, day *in vitro* (*DIV*) 9 cortical neurons cultured at high density (2.6 million in 100 mm culture dishes) were washed three times with HBSS. After the final wash, HBSS was replaced with 4 ml Neurobasal media per 100 mm dish. Neurobasal media does not contain serum but contains growth factors. Neurobasal conditioned media was collected after 4 h incubation and processed for differential ultracentrifugation. Briefly, conditioned media was centrifuged sequentially at 300xg for 10 min, 2000xg for 25 min in a tabletop centrifuge, and 10,000xg for 30min in a Type 45 Ti ultracentrifuge rotor (Beckman) to remove live cells, cell debris and LEVs, respectively. The supernatant from the 10,000xg spin was centrifuged at 100,000xg for 18 hrs in a Type 45 Ti rotor to obtain SEVs. The 100,000xg SEV-containing pellets were resuspended in 3 ml sterile cold PBS and repelleted at 100,000xg for 4 hrs in a TLA110 rotor (Beckman). SEVs were analyzed for size and number by nanoparticle tracking (ZetaView, ParticleMetrix), for morphology by TEM, and for common SEV markers by Western blotting.

#### Western Blot Analysis

Neuron TCL and EV samples for Western blot were prepared as described above in the cancer cell Western blot methods section. Antibodies (for neurons): rabbit anti-TSG101 (Abcam ab30871), mouse anti-Flotillin-1 (BD Biosciences Cat# 610820), Mouse anti-Alix (Cell Signaling Cat# 2171), mouse anti-GM130 (BD Biosciences Cat# 610822), rabbit anti-THSD7A (HPA000923, Sigma).

#### Calcium Phosphate Transfection

Neurons plated at low density on coated glass coverslips were transfected with a modified calcium phosphate transfection method at day 3 or 5 as previously described ^112^. 1 μg of mCherry-pTαS2 and either 1 μg of GFP-pTαS2 or 3 μgs of GFP-Rab27b/ shRNAs or 2 μgs of pCMV-FLAG/pCMV-FLAG-THSD7A were mixed with 120μl of 250mM CaCl_2_ in an Eppendorf tube. 120 μl of 2X HEPES Buffered Saline (HBS) (274 mM NaCl, 9.5 mM KCl, 15 mM glucose, 42 mM HEPES, 1.4 mM Na_2_HPO_4_, pH 7.15) was added drop by drop to the DNA-CaCl_2_ mixture with continuous aeration and incubated at room temperature for 15 min. The neuron coverslips were removed from co-cultures and transferred to sterile petri dishes, containing glial conditioned media, for dropwise addition of the transfection mixtures. Neurons were kept in the incubator at 37° C for 30-40 minutes and then washed three times with HBSS (135 mM NaCl, 4 mM KCl, 2 mM CaCl_2_, 1 mM MgCl_2_, 10 mM glucose, 20 mM HEPES, pH 7.35). The neuron coverslips were then transferred back to the home dishes containing astrocytes. The transfection efficiency was in the range of 5-10% using this method. Knockdown or expression of target genes was analyzed by immunofluorescence together with expression of co-transfected mCherry filler (shRNAs, see Fig. S4A-D) or fluorescence (GFP or GFP-Rab27a). Analysis of KD phenotypes was performed in transfected neurons, as assessed by fluorescence of the mCherry filler, as in previous publications ^113^.

#### Immunocytochemistry

For most antibodies, neurons were fixed in 4% paraformaldehyde in PBS for 15 min at RT and then permeabilized with 0.2% triton-X in PBS for 5 min at RT. For PSD95 staining, neurons were fixed in 4% paraformaldehyde in PBS for 3 min at RT and then incubated in cold methanol for 10 min at −20^°^C. Next neurons were incubated for 1 hour with 20% goat serum to block non-specific antibody binding. Primary antibody was diluted in 5% goat serum in PBS with 1:1000 dilutions of antibody. Neurons were incubated with the primary antibody mixture overnight at 4^0^C. The secondary antibody mixture was prepared similarly to the primary antibody mixture. Neurons were incubated for 1 hour at room temperature with the secondary antibody mixture. Cells were washed three times with 1x PBS between each step. After the final wash with PBS, neuron coverslips were mounted on glass slides using AquaPoly Mount and allowed to dry overnight.

#### Microscopy and Image Analysis

Neuronal imaging was performed on a Quorum Wave-FX Yokogava CSU-X1 spinning disk confocal system with a Nikon Eclipse Ti microscope. Images were acquired using MetaMorph software (Molecular Devices, Sunnyvale, CA) and a Plan Apo TIRF 60x (NA 1.49) objective. Images for GFP, mCherry and SV2 647/PSD95 647 were acquired by laser excitation at 491 nm, 561 nm and 642 nm respectively. Emission filters for these fluorophores were 525/50, 593/40 or 620/60 and 700/75 respectively (Semrock, Rochester, NY). Primary or secondary dendrites from confocal images were randomly selected for quantification of filopodia and spine density. Dendritic filopodia were defined as thin headless protrusions. Dendritic spines were identified as dendritic protrusions that co-localize with synaptic markers SV2. Synapses were defined as SV2-positive puncta present on both dendritic protrusions and dendritic shafts.

## Acknowledgments

Funding was provided by NIH R01GM117916, R01CA206458, R01CA249684, and R01CA249424 grants to AMW, by AHA fellowship 17POST33660473 and NIH training grant T32CA009592 support of COM, by Grant-in-Aid for JSPS KAKENHI (JP 17K15005) to DH, by NIH R50CA283661 to BHS. This project was begun as a collaborative project with Dr. Donna Webb, who passed away before her time. We acknowledge her important scientific contributions to this work, including her mentorship of Dr. Mikin Patel, her expertise in neuroscience and cell adhesion mechanisms, and her insightful suggestions.

---

**Supplemental Figure 1. Characterization of B16F1 Cells and EV Preparations.** A. Western blot of Hrs-KD in B16F1 cells. B. Western blot of Rab27a-KD in B16F1 cells. C. Nanoparticle tracking analysis representative traces showing particle size distributions that correspond to the data for control SEVs and LEVs in Fig. 2A. D. Western blot of gradient fractions for SEVs prepared by the cushion density gradient (DG) method. TCL, total cell lysate. LEVs also run on gel. E. TEM of negatively stained LEVs and SEVs prepared by the cushion-DG method. Scale bar = 200 nm in each image.

**Supplemental Figure 2. Characterization of HT1080 Cells and EV Preparations and comparison of graphing methods.** A. WB of Rab27a KD in HT1080 cell lysates. B. TEM of SEVs (purified by cushion DG) and LEVs from HT1080 cells. Scale bar = 200 nm in each image. C. Secretion rates of SEVs from HT1080 cell lines (N=3). Nanoparticle tracking analysis traces of SEVs from shScr and shRab27a HT1080 cells showing size (diameter) distribution of SEVs and particles/mL/cell. D. Representative images showing filopodia in control and Rab27a-KD H1080 cells. Images have been edited with brightness and contrast tools for ease of visibility. Scale bars in wide field and zoom insets = 10 μm. E. Quantification of filopodia in control and Rab27a-KD HT1080 cell lines. N=3, ≥ 20 cells per condition per rep. F. Data from graph in E displayed as filopodia per cell. G. Data in Figure 2C displayed as filopodia per cell. H. Data from Figure 2E displayed as filopodia per cell. I. Data from Figure 2F displayed as filopodia per cell. J. Data from figure 2G displayed as filopodia per cell. K. Cell areas of cells used for quantification in Figure 2G and Supp Figure 2J. Error bars, SEM. ns, not significant; * p<0.05; ** p<0.01; *** p<0.001.

**Supplemental Figure 3. Exosome secretion enhances filopodia, spine and synapse numbers in primary hippocampal neurons.** A. and B. Primary rat hippocampal neurons were co-transfected with GFP-Rab27b or GFP, and mCherry filler (red) on *DIV*5 and immunostained for SV2 (cyan) on *DIV*6. Overlay images showing localization of GFP-Rab27b at the base and tips of filopodia and quantification of percent GFP-Rab27b localization at tips and bases of filopodia in hippocampal neurons are shown in Fig. 1E,F. A. Representative mCherry and SV2 overlay images. B. Quantification of filopodia, spine and synapse density from images as in A. C-F. Hippocampal neurons were transfected with NTshRNA or shRNAs against Rab27b (C,D) or Hrs (E,F) and immunostained for SV2 at *DIV*6 for filopodia analysis or *DIV*12 for spine and synapse analysis. Quantification of number of filopodia, spines and synapses for Rab27b-KD (D) or Hrs-KD (F) neurons. Filopodia, spine and synapse density quantified from ≥ 30 primary or secondary dendritic shafts from three independent experiments for each condition. Scale bar = 5 μm. Error bars, SEM. *p<0.05, **p<0.01, ***p<0.001.

**Supplemental Figure 4. Analysis of exosome gene knockdown and SEV rescue in cortical neurons.** A. and B. Representative images of neurons co-transfected with NTshRNA, Hrs shRNA, or Rab27b shRNA, and mCerulean, as indicated, and immunostained for Hrs or Rab27b. Blue arrows indicate mCerulean-expressing neurons, which are the ones assumed to be cotransfected with shRNA and are quantitated for filopodia, spine and synapse density. White arrows indicate non-mCerulean-expressing neurons. C. and D. Quantitation of Hrs (C) or Rab27b (D) immunofluorescence levels in mCerulean-expressing neurons for the indicated transfection conditions. E-G, analysis of SEVs purified from cortical neurons (CN). E. Nanoparticle tracking analysis trace showing size distribution. F. Western blot analysis of positive (TSG101, Flotillin-1, Alix) and negative (GM130) markers. G. Negative stain TEM image. H. and I. Representative images of control and exosome-KD cortical neurons +/-addback of purified SEVs (+SEV). Arrows indicate filopodia. Scale bar = 5 μm. Quantitation is in Fig. 3I,J. Error bars, SEM. *p<0.05, **p<0.01, ***p<0.001.

**Supplemental Figure 5. KD of Endoglin affects filopodia numbers but not EV numbers in B16F1 melanoma cells.** A. Nanoparticle tracking analysis traces for B16F1 control (shScr) and endoglin-KD (shEng) SEVs. B. SEV secretion rates from B16F1 shEng stable lines. N=5. C. Representative Western blot of Endoglin-KD in transient siRNA-transfected B16F1 cells. D. Filopodia numbers in siRNA-transfected B16F1 cells (N=3, ≥ 23 cells per condition per rep). E. Images from control and shHrs cells incubated with purified EV, corresponding to graph in Fig. 4F. Scale bar in wide field and zoom insets = 10 μm. Error bars, SEM. ns, not significant; * p<0.05; ** p<0.01; *** p<0.001.

**Supplemental Figure 6. HT1080 Endoglin-KD cells have reduced filopodia.** A. Western blot of Endoglin KD in HT1080 cells. B. Nanoparticle tracking analysis traces of SEVs purified from shScr and shEng HT1080 cells showing size distribution (diameter) of SEVs and particles/mL/cell (N=3). C. SEV secretion rates of HT1080 shScr, and shEng HT1080 cells. D. Representative images of HT1080 shScr and shEng cells. Images have been edited with brightness and contrast for ease of visibility. Scale bar in wide field and zoom insets = 10 μm. E. Quantitation of filopodia density for control and shEng HT1080 cells. N=4, 20 cells per condition per rep. Error bars, SEM. ns, not significant; * p<0.05; ** p<0.01; *** p<0.001.

**Supplemental Figure 7. Analysis of candidate EV cargoes for rescue of filopodia defects in shEng cells.** A. Western blot showing β1-integrin, TGFβ1, ALK1 levels in control (shScr) and endoglin-KD (shEng) B16F1 SEVs. B. Filopodia density analysis of B16F1 shScr and shEng cells treated with BMP-9. N=3, ≥ 20 cells per condition per rep. C. Filopodia density analysis of B16F1 shScr and shEng cells treated with TGFβ1. N=3, ≥ 20 cells per condition per rep. D. Filopodia density analysis of B16F1 shScr and shEng cells plated on 20 µg/ml fibronectin (FN) or 100 µg/mL poly-D-lysine (PDL). N=3, ≥ 20 cells per condition per rep. E. Filopodia density analysis of B16F1 shScr and shEng cells plated on PDL or 2 µg/mL rhTHSD7A for the indicated time points, then fixed and stained for filopodia. N=3, at least 20 cells per condition per rep. F. Total cell lysates of B16F1 cells were seeded on PDL +/-THSD7A and treated with or without TGF-β1 and inhibitor. Cell lysates were probed for phospho-Smad and total Smad levels. Representative of 3 blots. Error bars, SEM. ns, not significant; * p<0.05; ** p<0.01; *** p<0.001.

## Supplemental Movies

**Supplemental Movie 1. Visualization of exosome secretion and filopodia formation dynamics.** HT1080 cells stably expressing pHLuorin_M153R-CD63-mScarlet were seeded onto glass bottom MatTek plates and imaged live. Images were taken every 10 seconds.

**Supplemental Movie 2. Live imaging of filopodia in B16F1 control and shHrs cells.** B16F1 cells stably expressing shHrs or control plasmid (shLacZ) were transiently transfected with tdTomato-fTractin and seeded onto PDL-coated glass bottom MatTek Plates 24 hours post-transfection. Live cells were imaged every 30 seconds for 15 minutes. Movies were inverted, de-noised, and brightness/contrast was adjusted to facilitate visualization of filopodia.

**Supplemental Movie 3. Live imaging of filopodia in B16F1 control and shEng cells.** B16F1 cells stably expressing shEng or control plasmid (shScr) were transiently transfected with tdTomato-fTractin and seeded onto PDL-coated glass bottom MatTek plates 24 hours post-transfection. Live cells were imaged every 30 seconds for 15 minutes. Movies were inverted and brightness/contrast was adjusted to facilitate visualization of filopodia.

**Supplemental Movie 4. Endoglin controls 3D migration in type I collagen.** The indicated HT1080 cell types were induced to form spheroids and then mixed into 3D type I collagen.

Spheroids were imaged every 30 minutes for 8 hours. Scale bar = 100 μm.

## Supplemental Tables

Supplemental Table 1: Proteomics results from bands cut from colloidal blue gel in Figure 4A.

Supplemental Table 2: iTRAQ proteomics results from control and endoglin-KD B16F1 SEVs.

## Notes

### Competing Interest Statement

The authors have declared no competing interest.

### Summary of Updates

We made some text changes to clarify specific points and include new references and discussion. We also edited the figures to add scale bars and labels, and to add a comparison of quantitating filopodia/cell area to filopodia/cell in Supplemental Figure 2. We also split Figure 8 into two figures so that the model (formerly Fig 8b,c) would have its own figure (Fig 9).

