## Supplementary figures and images for "Secreted exosomes induce filopodia formation"

### Supplementary Figures

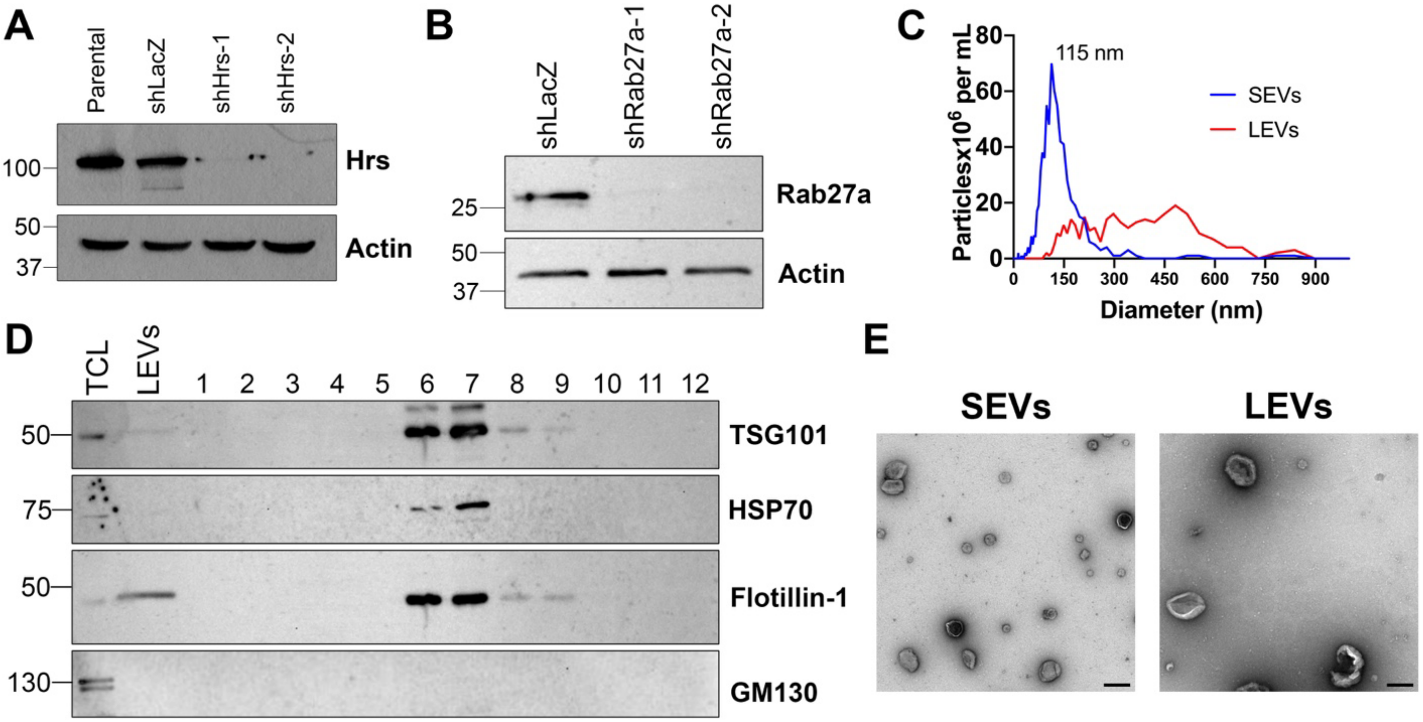

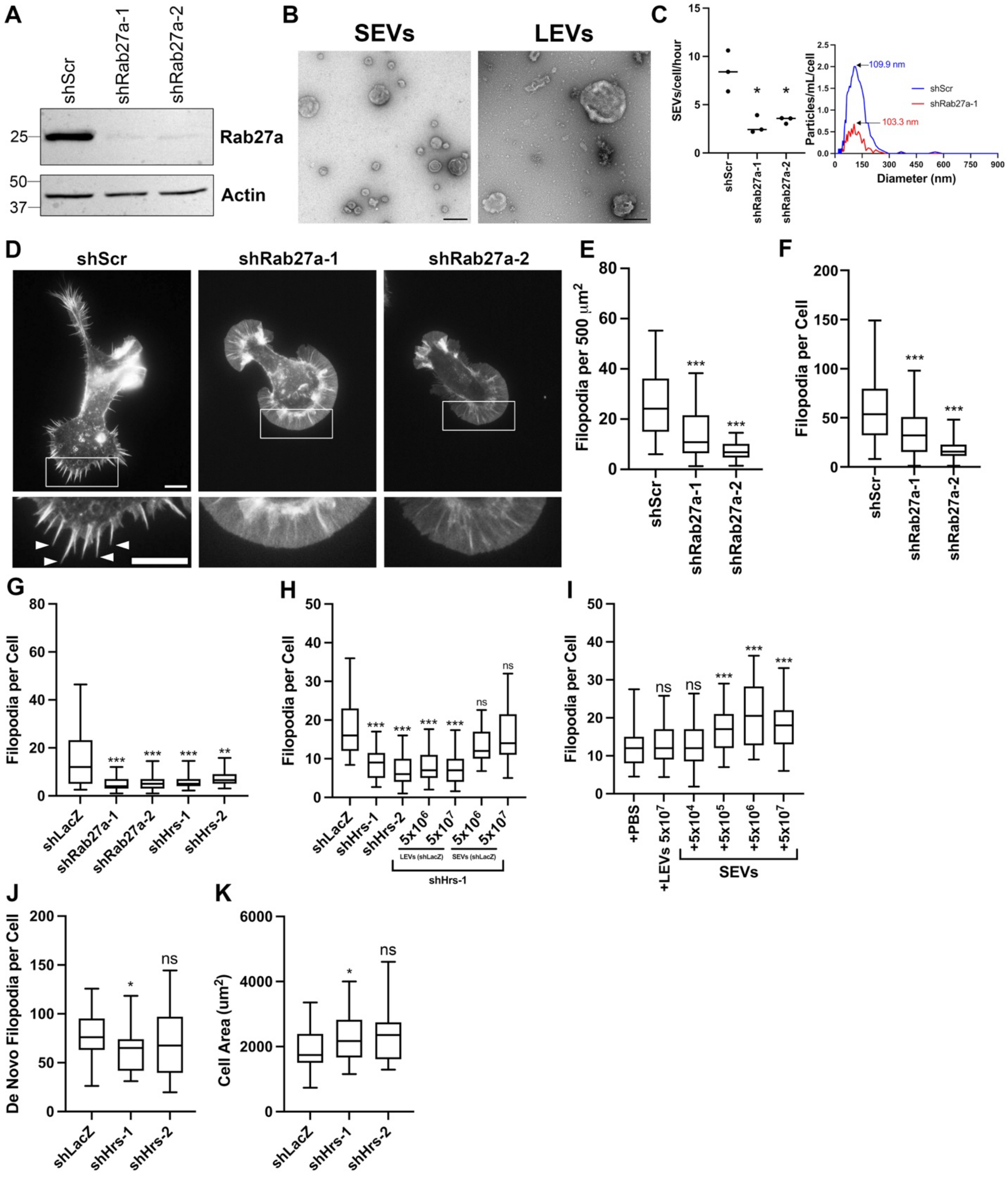

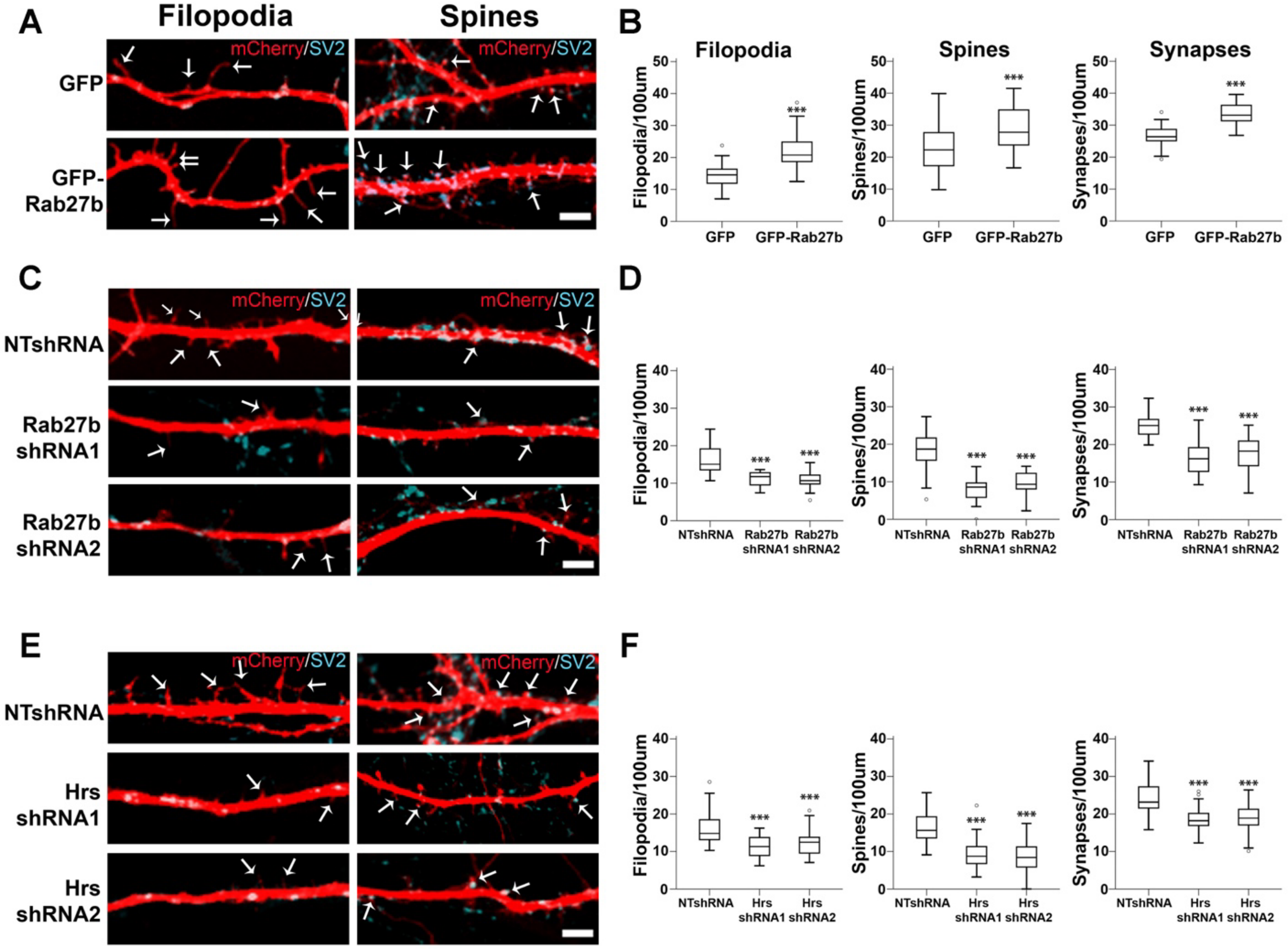

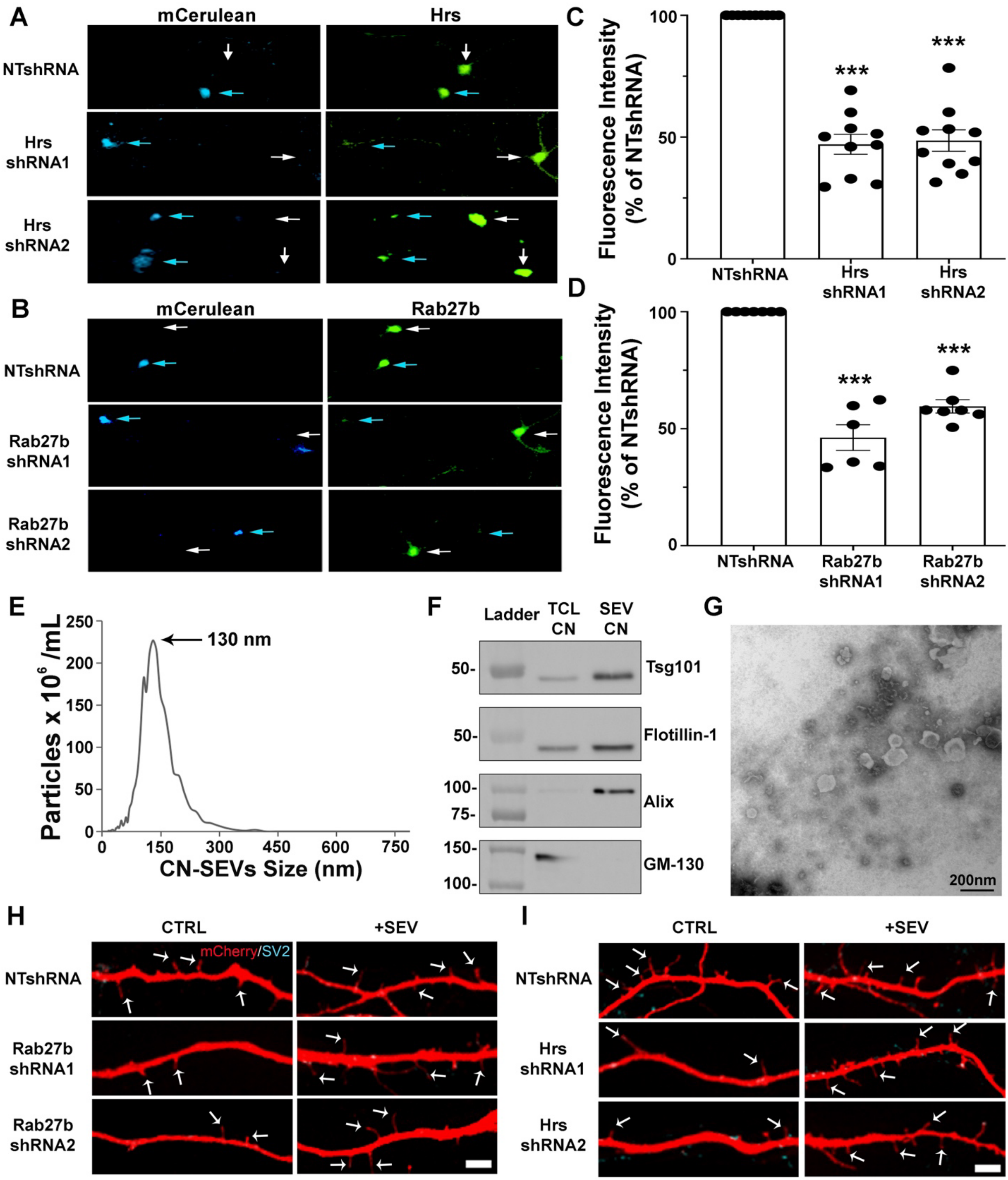

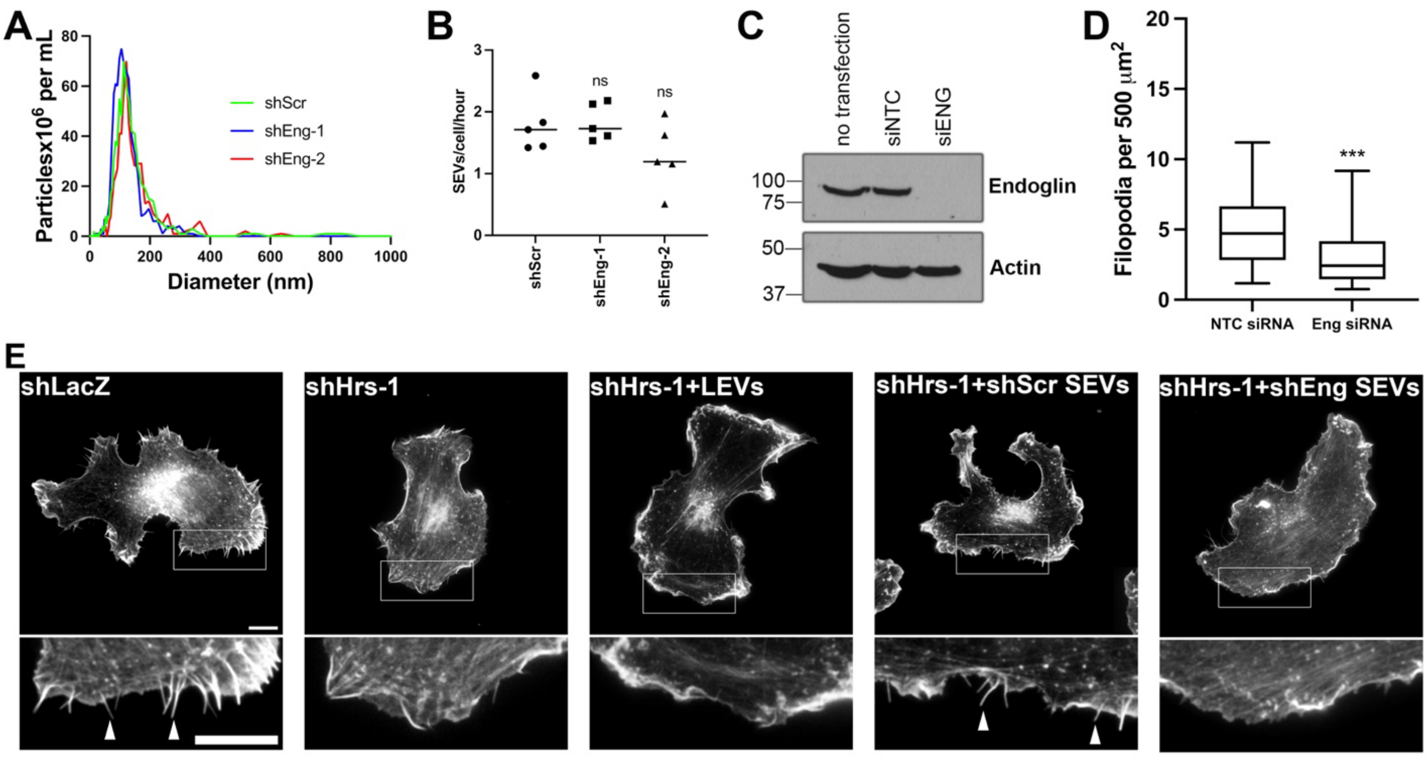

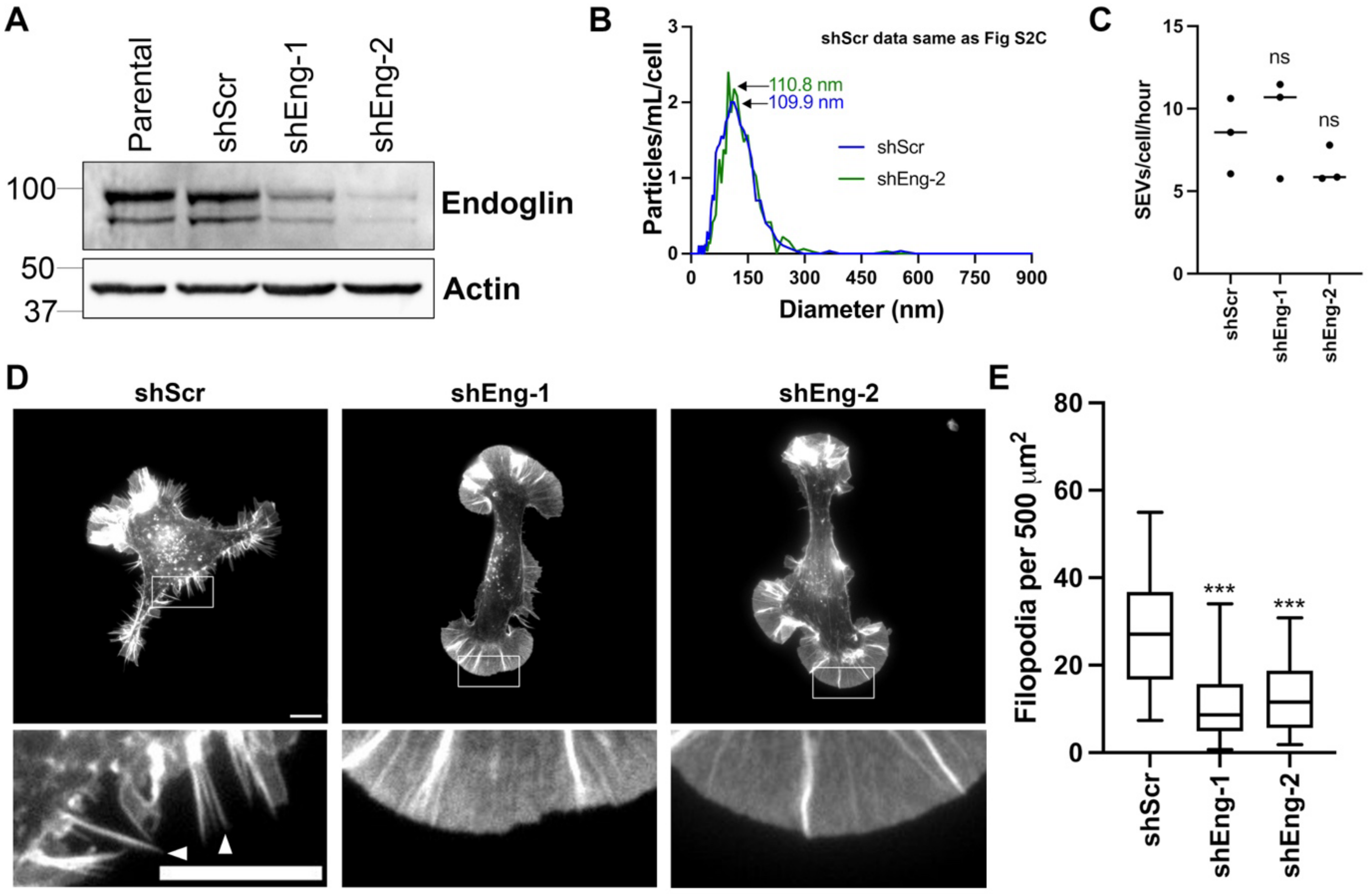

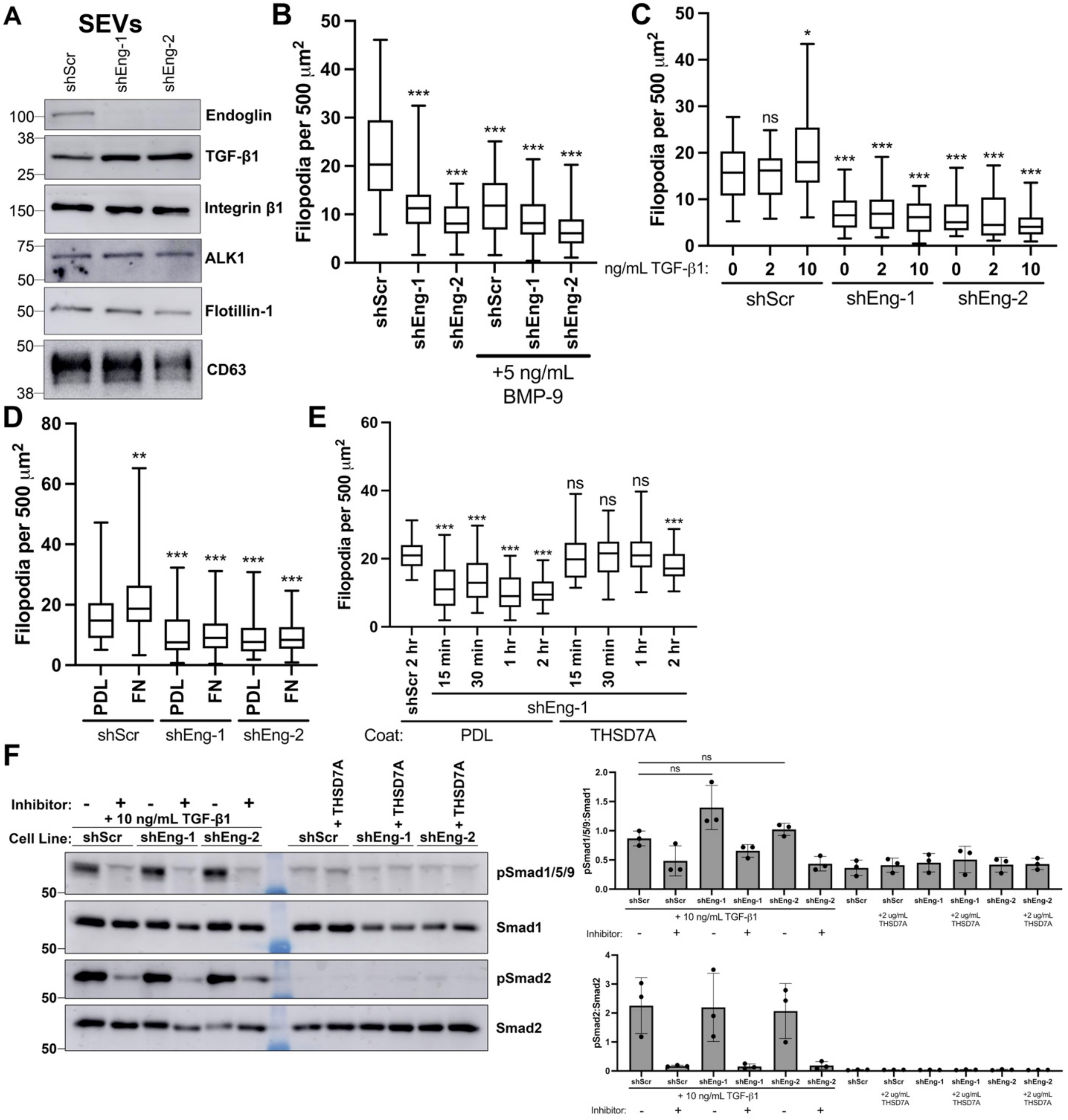
